# SAD-6/ATRX enables broad genome surveillance and defense in fungi

**DOI:** 10.64898/2026.08.11.744009

**Authors:** Florian Carlier, Anna Klimova, Eleonore Bouscasse, Ziyan Wang, Isabelle Loïodice, Angela Taddei, Ilkka Kronholm, Jay C. Dunlap, Mariette Matondo, Eugene Gladyshev

## Abstract

The chromatin remodeler ATRX and its orthologs maintain genome function by regulating repetitive DNA and dynamic chromatin, and their activities have been canonically associated with replication-independent deposition of the histone H3.3 variant. This model is difficult to reconcile with fungi, which encode ATRX orthologs but lack H3 variants that may separately support replication-coupled and replication-independent deposition. Here we show that the fungal ATRX ortholog SAD-6 instead relies on a highly divergent histone H4 variant (H4v) to mediate broad genome surveillance and defense. Deposition of H4v is strictly SAD-6-dependent and thus provides a sensitive genome-wide readout of SAD-6 activity, revealing its functions at telomeres, tRNA and rDNA loci, AT-rich DNA, artificial transgenes, decaying mobile elements, and many genic regions. We further show that SAD-6 is required for a pathway of repeat-induced point mutation (RIP) that also requires DIM-5, a conserved SUV39 methyltransferase that mediates trimethylation of histone H3 lysine-9 in heterochromatin. Together, these findings establish ATRX-like remodelers as broad regulators of genome surveillance and defense in fungi that act through a highly divergent histone H4 variant rather than H3.3. Given that RIP is proposed to recognize repetitive DNA via recombination-independent homologous pairing, the requirement for SAD-6 in RIP suggests that ATRX-like remodelers may couple DNA pairing to heterochromatin nucleation on repeats.

## INTRODUCTION

ATRX (alpha-thalassemia/mental retardation syndrome X-linked) was first identified through genetic studies of patients with ATR-X syndrome, a neurodevelopmental disorder associated with intellectual disability and alpha-thalassemia^1^. Mammalian ATRX carries an N-terminal ADD (ATRX-DNMT3-DNMT3L) domain and a C-terminal SF2 helicase domain; and together with the histone chaperone DAXX (death-domain associated protein), it mediates replication-independent deposition of the histone variant H3.3 at diverse types of DNA repeats, such as dispersed copies of transposable elements, ribosomal DNA, pericentromeres, and telomeres^2^. This process is frequently coupled to heterochromatin assembly, thereby suppressing aberrant recombination and limiting replication stress^2^. In parallel, ATRX can also be recruited to actively transcribed genes by a yet unknown mechanism that appears to be *process*-dependent rather than sequence-dependent^3^. ATRX orthologs are broadly conserved across eukaryotes^4–7^, where they also mediate H3.3 deposition in dynamic chromatin.

The fungus *Neurospora crassa* encodes a single ATRX ortholog, SAD-6 (suppressor of ascus dominance-6), which was originally implicated in meiotic silencing by unpaired DNA (MSUD)^8^. Subsequently, SAD-6 was found to control heterochromatin assembly and siRNA production from chromatin perturbed by the aberrant binding of non-histone proteins^9^. In that system, effectively the same silencing response was induced by two different stimuli (binding of a synthetic bacterial repressor TetR-GFP or the endogenous transcription factor NIT-2)^9^, both associated with reduced nucleosome occupancy and elevated MNase sensitivity, two hallmark features of ATRX targets in animals and plants^9^. However, like many other fungi, *N. crassa* has only two H3 histones, one canonical and one centromeric^10^, suggesting that fungi may provide an exception to the H3.3-centered function of ATRX.

In this study, we investigated the function of SAD-6 as a model ATRX ortholog in fungi. We first developed a forward-genetic strategy to identify factors required for silencing of a repressor-bound *tetO* array carrying a selectable marker. This approach uncovered the histone H4 variant H4v (NCU04338) as an essential cofactor for SAD-6-dependent heterochromatin formation and siRNA production. Genome-wide ChIP-seq profiling identified numerous H4v-enriched regions, including the *tetO* array, tRNA genes, rDNA loci, telomeric and AT-rich regions, decaying mobile elements, as well as intergenic and gene-proximal regions. H4v deposition required SAD-6 in every context, providing a sensitive genome-wide readout of SAD-6 activity. Several H4v peaks occurred at loci previously shown to produce Dicer-independent small interfering RNAs (disiRNAs) and undergo dynamic H3K9me3 and DNA methylation (5mC), including the circadian clock gene *frequency* (*frq*)^11–13^. Consistent with earlier work^12–14^, the transcription factor WC-2 was required for H4v enrichment at the *frq* promoter. Although H3K9me3 and 5mC at *frq* were lost in the absence of SAD-6, core clock function remained unaffected, suggesting that these marks may represent a downstream consequence of transcription-coupled chromatin remodeling rather than feedback regulation of the circadian clock.

We also examined the role of SAD-6 and H4v in repeat-induced point mutation (RIP), a premeiotic process that detects duplicated sequences through recombination-independent homologous DNA pairing and mutates them by introducing numerous C-to-T transitions^15^. RIP operates through two largely independent pathways: one mediated by the cytosine methyltransferase RID^16^ and another mediated by the methyltransferase DIM-2 downstream of DIM-5-dependent H3K9me3 deposition^17,18^. Notably, DIM-5 belongs to the SUV39 family of lysine methyltransferases that generally mediate H3K9me3 on repetitive DNA in eukaryotes^19^. We found that SAD-6 is essential for DIM-2-dependent RIP but dispensable for RID-dependent RIP, thereby identifying it as a critical component of the heterochromatin-related pathway of RIP.

Our findings give rise to four principal conclusions. First, ATRX-like proteins emerge as broad regulators of genome surveillance and defense in fungi. Second, the function of ATRX-like proteins does not universally require dedicated H3.3 variants. Third, in fungi, a highly divergent histone H4 variant fulfills an analogous genome-wide role. Finally, ATRX-like remodelers may serve as a link between recombination-independent homologous DNA pairing and heterochromatin nucleation on repetitive DNA.

## RESULTS

### Developing a forward genetic strategy to identify suppressors of SAD-6-dependent silencing

Reporter strain T838.4 was engineered to carry a *his-3*-proximal *tetO* array with a nourseothricin resistance gene (*ntcR*) and a TetR-GFP expression construct in place of *csr-1+* (Fig. 1A). Binding of TetR-GFP to the *tetO* array induced strong SAD-6-dependent silencing, repressing *ntcR* and rendering the strain sensitive to nourseothricin (NTC) (Fig. 1D,G; Fig. S1A). NTC resistance could be restored by deleting *dim-5+*, *dim-2+*, or *sad-6+* (Fig. 1D). Suppressors of silencing could thus be obtained by selection on NTC. Because loss of H3K9me3 in *N. crassa* triggers genome-wide redistribution of H3K27me3, which impairs growth and could potentially sustain silencing of *ntcR*, the reporter strain was constructed in a *set-7Δ* background, lacking the capacity to form H3K27me3^20^.

**Figure 1.**
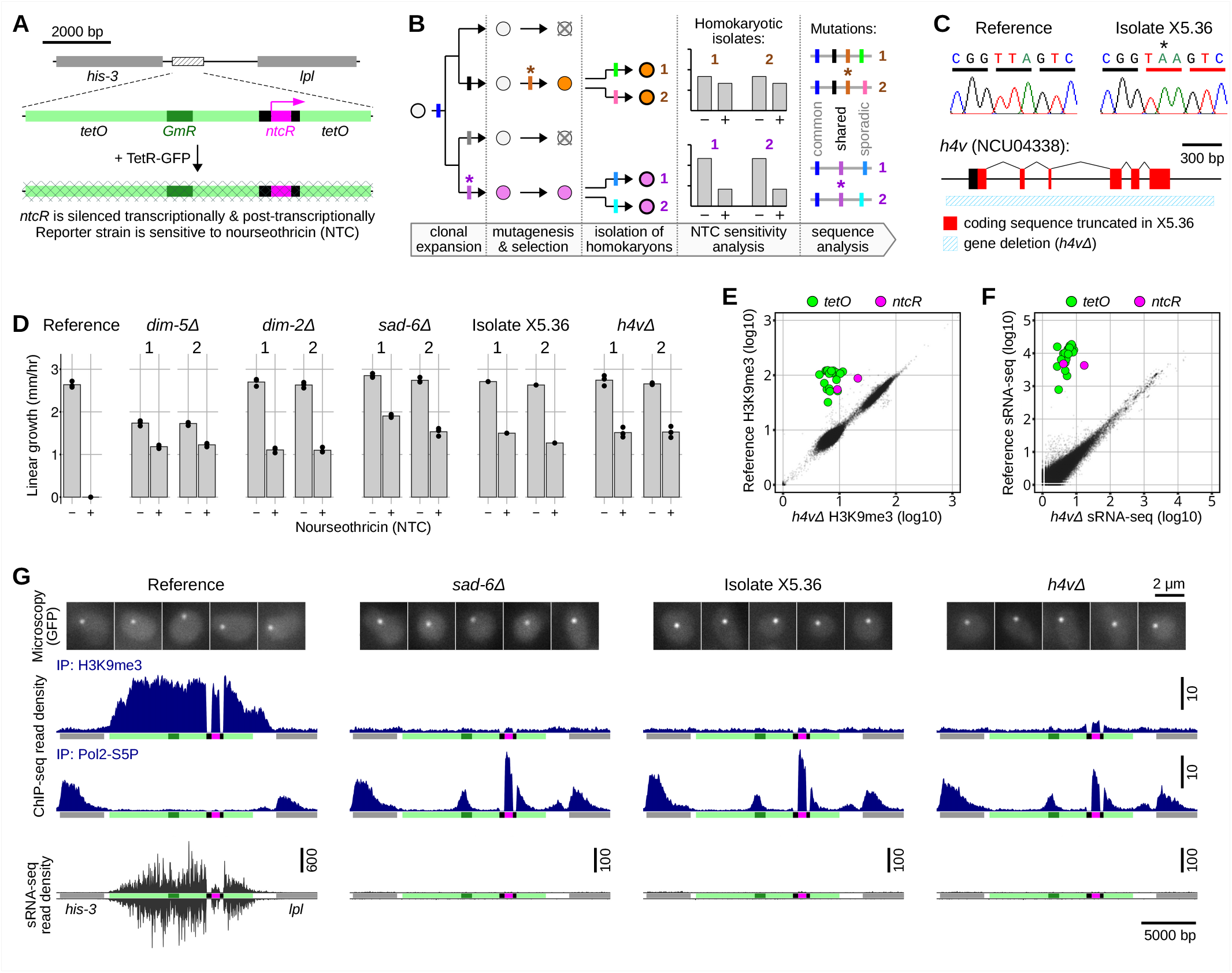
Histone H4 variant (H4v) is required for transcriptional and post-transcriptional silencing of the repressor-bound *tetO* array. **(A)** Schematic of the forward-genetic selection system. A *tetO* array with a nourseothricin-resistance gene (*ntcR*) was integrated between *his-3* and *lpl* in strain C139.10 to create strain T830.14h; a synthetic construct for constitutive expression of TetR-GFP was integrated in place of *csr-1+* in strain T830.14h to create strain T838.4 (this strain was also constructed in a *set-7Δ* background, lacking the capacity to form H3K27me3). Binding of TetR-GFP to the *tetO* array results in SAD-6-dependent silencing of *ntcR*, rendering the reporter strain T838.4 sensitive to nourseothricin (NTC). Black rectangles around *ntcR* correspond to regions masked due to their repetitive nature in the genome. Corresponding genome reference is provided in Dataset 1. Annotated sequences of transformation plasmids are provided in Dataset 2. **(B)** Strategy to identify suppressors of SAD-6-dependent silencing of *tetO::ntcR*. Macroconidia were plated on sorbose agar containing NTC and UV-mutagenized at low dose. NTC-resistant colonies were propagated via uninucleate microconidia to establish homokaryotic isolates, which were then assayed for linear growth in the presence/absence of NTC and subjected to whole-genome sequencing. SNPs uniquely shared by a pair of homokaryotic isolates originating from the same primary NTC-resistant colony were considered candidate causal mutations. Asterisks mark hypothetical mutations that confer resistance to NTC. **(C)** Identification of a mutation in the histone H4 variant gene *h4v* (NCU04338) in isolate X5.36. A T-to-A substitution introduces a premature stop codon. A corresponding deletion allele (*h4vΔ*) is shown below. **(D)** Linear growth of *N. crassa* strains in the presence/absence of NTC. Deletion of *dim-5+*, *dim-2+*, *sad-6+,* or *h4v+* restores growth on NTC, indicating a defect in *tetO::ntcR* silencing. Two independent transformants were analyzed for each condition: T914.1h and T914.3h (*dim-5Δ*); T923.1h and T923.2h (*dim-2Δ*); T902.1h and T902.4h (*sad-6Δ*); X5.36.M1 and X5.36.M2 (X5.36 isolate); T922.4h and T922.6h (*h4vΔ*). Reference condition corresponds to strain T838.4. **(E)** Genome-wide comparison of H3K9me3 ChIP-seq profiles between the reference and *h4vΔ* backgrounds. Signal was computed as the number of reads in 500-bp bins, normalized by the total coverage equivalent of one million reads, and a pseudo-count of 1 was added prior to log10 transformation. The following strains were analyzed: T838.4 (Reference); T922.1h and T922.6h (*h4vΔ*). **(F)** Genome-wide comparison of sRNA production between the reference and *h4vΔ* backgrounds. Signal was computed as the number of reads in 500-bp bins, normalized by the total number of aligned reads, a pseudo-count of 1 was added before log10 transformation. The following strains were analyzed: T838.4 (Reference); T922.4h and T922.6h (*h4vΔ*). **(G)** TetR-GFP localization and molecular profiles at the *tetO::ntcR* locus in several genetic backgrounds, as indicated. Top: representative fluorescence microscopy images on nuclei (n = 5). Below: normalized ChIP-seq (H3K9me3 and Pol2-S5P) and small RNA-seq profiles. The analyzed strains were as follows. ChIP-seq: T838.4 (Reference); T902.1h and T902.4h (*sad-6Δ*), X5.36.M1 and X5.36.M2 (isolate X5.36), T922.1h and T922.6h (*h4vΔ*). sRNA-seq: T838.4 (Reference); T902.1h and T902.4h (*sad-6Δ*), X5.36.M1 and X5.36.M2 (isolate X5.36), T922.4h and T922.6h (*h4vΔ*). Fluorescence microscopy: T838.4 (Reference); T902.1h (*sad-6Δ*), X5.36.M1 (isolate X5.36), T922.1h (*h4vΔ*). Scale bar, 2 µm.

Suppressor mutations were identified as follows (Fig. 1B). A clonal batch of macroconidia (∼5 × 10⁸ CFUs, derived from a single uninucleate microconidium) was first plated directly on NTC-containing medium and UV-mutagenized at a low dose. Because macroconidia are multinucleate, each primary NTC-resistant colony was propagated through microconidia to produce several homokaryotic isolates, which were then assayed for linear growth in the presence/absence of NTC and subjected to whole-genome sequencing. Candidate causal mutations were identified as single-nucleotide polymorphisms (SNPs) uniquely shared by homokaryotic isolates originating from the same primary NTC-resistant colony. Because mutagenesis was performed at a low dose, only a small number of SNPs had to be considered per primary isolate.

### Histone H4 variant (H4v) is required for SAD-6-dependent silencing of the *tetO::ntcR* reporter

One NTC-resistant isolate (X5.36) contained a nonsense T-to-A mutation predicted to truncate the histone H4 variant H4v (NCU04338)^21^ (Fig. 1C,D). The causative nature of this mutation was confirmed by deletion of the entire *h4v+* gene (Fig. 1C,D,G). Overall, the *h4vΔ* and *sad-6Δ* strains shared three key characteristics: (i) normal growth in the absence of NTC, suggesting that loss of either factor does not broadly impair fitness in vegetative culture (Fig. 1D); (ii) robust growth in the presence of NTC, reflecting complete loss of silencing of the perturbed *tetO* array (Fig. 1D,G); and (iii) largely unchanged genome-wide levels of H3K9me3, Pol2-S5P (RNA polymerase II phosphorylated at serine-5 of the C-terminal domain), and small RNAs (Fig. 1E,F; Fig. S1B-D). These results place H4v and SAD-6 in the same silencing pathway and indicate that their loss specifically disrupts repression of a severely perturbed chromatin site (the *tetO* array) without causing global changes in heterochromatin, transcription, or small RNA production. Thus, H4v and SAD-6 together define a dedicated process that responds to perturbed chromatin.

### H4v is incorporated into MNase-resistant nucleosomes

*N. crassa* encodes three histone H4 proteins: two canonical ones with the identical amino-acid sequences and H4v, which is also broadly conserved in other filamentous fungi^22^. Although H4 and H4v share 62% identity, their sequences can be aligned without gaps (Fig. S2A). H4v contains a long N-terminal extension (25 amino acids) and a unique C-terminal part (35 amino acids) predicted to protrude from the nucleosome and form a novel structural element (Fig. 2A; Fig. S2A). C-terminally tagged versions of H4v differed in their effects on silencing: H4v-GFP was associated with a minor silencing defect, whereas no such effect was observed with H4v-FLAG (Fig. S2B). H4v-FLAG was therefore used for molecular analyses. The corresponding reference strains (T961 series) retained the *tetO* array but lacked both *ntcR* and *tetR-gfp*. In these strains, withdrawal of all nitrogen during the final 5 hours of liquid culture was used to induce acute nitrogen starvation and trigger SAD-6-dependent chromatin remodeling and silencing of *tetO*^9^.

**Figure 2.**
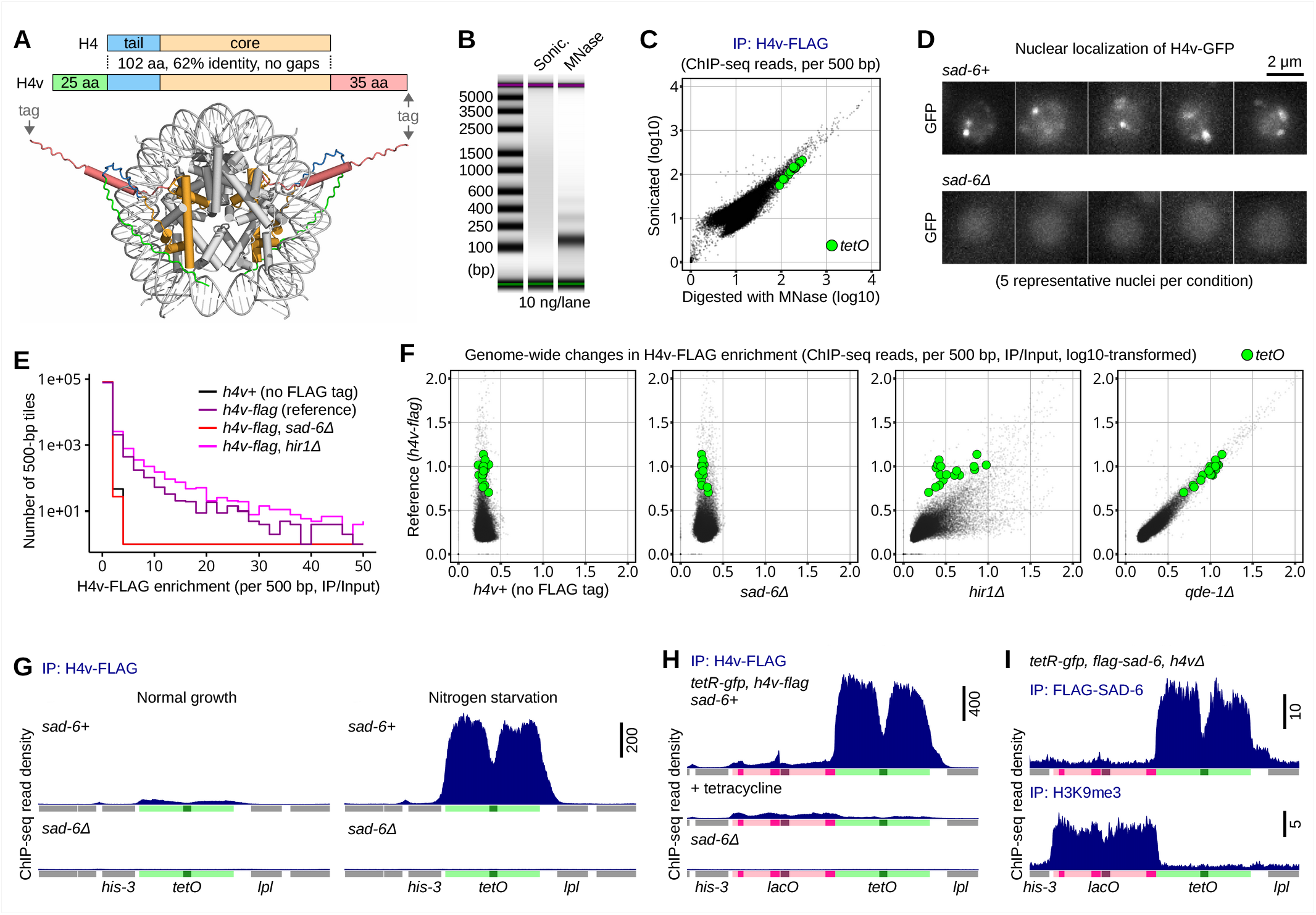
H4v chromatin enrichment requires SAD-6. **(A)** Comparison of the variant histone H4v and the canonical histone H4 of *N. crassa*. H4v contains an N-terminal extension and a unique C-terminal part predicted to form a novel structural element. A nucleosome with two H4v subunits modeled using the AlphaFold3 server (https://alphafoldserver.com) and visualized with PyMOL (Schrödinger) is shown. Five alternative predicted structures are provided in Dataset 3. **(B)** DNA fragmentation profiles generated by sonication versus MNase digestion. **(C)** Genome-wide comparison of H4v-FLAG ChIP-seq profiles originating from sonicated versus MNase-digested chromatin, analyzed and plotted as in Fig. 1E. The following strains were analyzed: T961.5h and T961.7h. **(D)** Nuclear localization of H4v-GFP. Representative fluorescence microscopy images of nuclei (n = 5) from wild-type and *sad-6Δ* strains expressing H4v-GFP are shown. The following strains were analyzed: T959.1h (*sad-6+*) and T964.1h (*sad-6Δ*). Scale bar, 2 µm. **(E)** Histogram showing genome-wide distribution of H4v-FLAG enrichment in several genetic backgrounds, as indicated. Signal was computed in 500-bp bins as a ratio of read counts normalized by the total coverage equivalent of one million reads. Bins with abnormally low Input values were excluded. The following strains were analyzed: T952.12h (untagged *h4v+*); T961.5h and T961.7h (*h4v-flag*); T965.4h and T965.9h (*sad-6Δ*); T968.1h and T968.3h (*hir1Δ*). **(F)** Genome-wide comparison of H4v-FLAG enrichment between several genetic backgrounds, as indicated. Signal was computed as in Fig. 2E, a pseudo-count of 1 was added prior to log10 transformation. The following strains were analyzed: T972.1h and T972.2h (*qde-1Δ*); the other analyzed strains are listed in Fig. 2E. **(G)** H4v-FLAG ChIP-seq profiles at the *tetO* array in the indicated conditions (normal growth versus nitrogen starvation; *sad-6+* versus *sad-6Δ*). The following strains were analyzed: T961.5h and T961.7h (*sad-6+*); T965.4h and T965.9h (*sad-6Δ*). **(H)** H4v-FLAG ChIP-seq profiles at the *lacO-tetO* tandem array in the indicated conditions. The following strains were analyzed: T948.1h and T948.7h (*sad-6+*); T949.1h and T949.2h (*sad-6Δ*). **(I)** FLAG-SAD-6 and H3K9me3 ChIP-seq profiles at the *lacO-tetO* tandem array in the *h4vΔ* condition. The following strains were analyzed: T950.4h and T950.6h.

We first asked whether H4v associates with chromatin and, more specifically, with nucleosomes. Chromatin from the reference strains carrying *h4v-flag* was fragmented by sonication or MNase digestion (Fig. 2B) and immunoprecipitated with an anti-FLAG antibody. Sonicated chromatin from a strain carrying untagged *h4v+* provided a negative control. Initial analysis of sonication-based ChIP-seq data revealed clear H4v enrichment over numerous 500-bp bins (Fig. 2E). Critically, ChIP-seq profiles generated from sonicated versus MNase-digested chromatin were highly concordant (Fig. 2C), even in MNase-sensitive regions that appear to contain nucleosomes in only a subset of nuclei (Fig. S2C). These results indicate that chromatin-bound H4v is likely found in nucleosomes, which are sufficiently stable to resist MNase digestion.

### H4v chromatin enrichment requires SAD-6

Using sonication-based ChIP-seq, we found that deletion of *sad-6+* caused a complete loss of H4v chromatin signal across the genome, including at the perturbed *tetO* array, which normally exhibits the highest levels of H4v (Fig. 2E-G). Consistent with the ChIP-seq data, loss of SAD-6 transformed the nuclear H4v-GFP signal from bright and heterogeneous to faint and uniform (Fig. 2D). Notably, the punctate appearance of H4v-GFP foci in *sad-6+* nuclei suggests that a fraction of H4v may reside in discrete subnuclear compartments.

### H4v chromatin enrichment is influenced by HIRA

The histone chaperone HIRA provides an alternative route for replication-independent deposition of H3–H4 dimers, acting in parallel with the SAD-6/ATRX pathway^23^. We found that deletion of *hir1+*, which encodes a core subunit of the HIRA complex, caused genome-wide changes in H4v enrichment, with an overall trend toward increased H4v occupancy (Fig. 2E,F). Thus, HIRA appears to limit H4v accumulation, either directly or indirectly. A direct effect could occur if HIRA-mediated deposition of canonical H3–H4 dimers competes with SAD-6-dependent deposition or retention of H4v-containing nucleosomes, and so loss of HIRA allows H4v to accumulate in chromatin. An indirect effect could occur if impaired nucleosome assembly in *hir1Δ* strains increases the abundance of nucleosome-depleted sites, thus expanding the pool of substrates available for SAD-6-dependent H4v deposition. In contrast, loss of the RNA-dependent RNA polymerase QDE-1 (a core component of vegetative RNAi) had little effect on H4v enrichment (Fig. 2F), consistent with a notion that chromatin organization is largely independent of RNAi in this organism^17^.

### SAD-6 is recruited to perturbed chromatin independently of H4v

Having found that SAD-6 is required for H4v association with chromatin, we next tested whether H4v is, in turn, required for SAD-6 recruitment to perturbed chromatin. Here we used a previously established system, in which FLAG-tagged SAD-6 is expressed from its native locus under the control of an inducible promoter^9^. This system is based on strain T658.1, which carries the *tetO* array integrated together with a similarly sized *lacO* array and also has *tetR-gfp* in place of *csr-1+*^9^. Whereas H3K9me3 at the repressor-bound *tetO* array is completely dependent on SAD-6, the *lacO* array exhibits constitutive SAD-6-independent H3K9me3, likely directed by AT-rich motifs within the *lacO* sequence^9^. Using this system, we previously reported that FLAG-SAD-6 is recruited specifically to the repressor-bound (perturbed) *tetO* array^9^.

Expressing H4v-FLAG in T658.1, we found that H4v was highly enriched at the repressor-bound *tetO* array, as expected (Fig. 2H). H4v enrichment was nearly abolished by displacing TetR-GFP using tetracycline, and it was completely eliminated in *sad-6Δ* strains (Fig. 2H), indicating that the initiating chromatin perturbation and SAD-6 are both required for H4v localization in this system. Interestingly, H4v was also observed at the *lacO* array, although at substantially lower levels than at the *tetO* array (Fig. 2H). However, deletion of *h4v+* did not eliminate H3K9me3 at *lacO* (Fig. 2I), suggesting that low-level H4v enrichment at this constitutively heterochromatic locus is not required for its maintenance.

Conversely, deletion of *h4v+* abolished H3K9me3 at the *tetO* array while leaving FLAG-SAD-6 occupancy unchanged (Fig. 2I; compare to Fig. 2G in ref. 9). These results separate SAD-6 recruitment from the downstream response, indicating that SAD-6 localizes to perturbed chromatin independently of H4v but requires H4v to mediate subsequent silencing.

### H4v forms a specialized complex with H3 and ASF1

Although the animal ATRX–DAXX–H3.3 system is considered prototypical, DAXX is absent in plants^7^, and both DAXX and H3.3 analogs are absent in fungi, suggesting that ATRX orthologs in these lineages function through mechanisms independent of the DAXX–H3.3 pathway. To identify histone chaperones that associate with H4v in *N. crassa*, sonicated whole-cell lysates from a strain expressing H4v-FLAG were fractionated by gel filtration chromatography on a Superdex-200 column and the fractions were assayed by Western blotting (Fig. 3A). H4v was found in fractions 2-3, which contained high-molecular-weight material, and in fractions 8-9, which eluted between β-Amylase and bovine serum albumin (BSA) with molecular masses of ∼200 kDa and ∼66 kDa, respectively (Fig. 3A). Based on column calibration, the putative H4v-containing complex in fractions 8-9 had an apparent molecular mass of ∼124 kDa, which may not directly reflect its stoichiometry.

**Figure 3.**
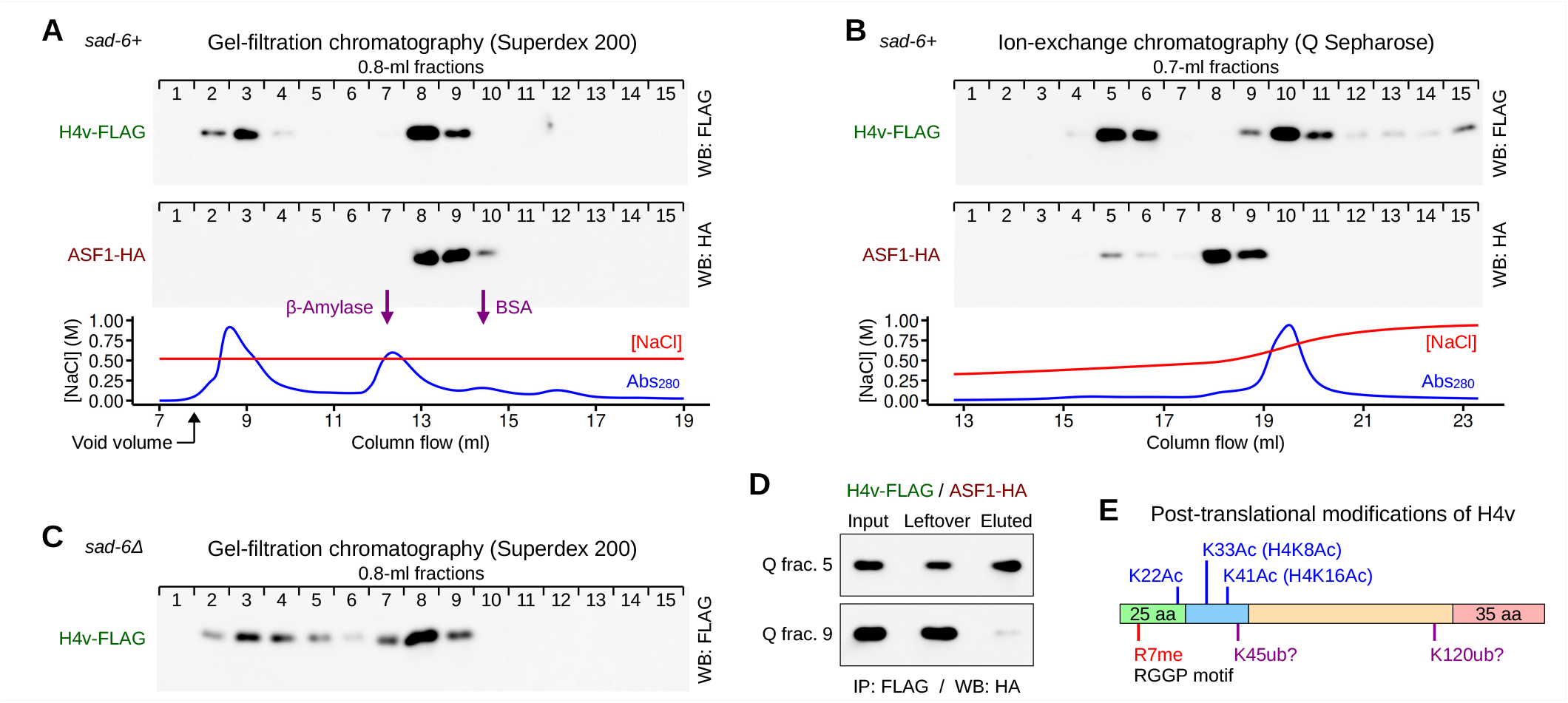
H4v forms a complex with ASF1 and H3. **(A)** Gel filtration chromatography (Superdex 200) of whole-cell lysates from strains expressing H4v-FLAG and ASF1-HA. Fractions were analyzed by Western blotting using anti-FLAG and anti-HA antibodies. At least two independent experiments were performed; a representative result is shown. H4v-FLAG co-elutes with ASF1-HA in fractions 8-9, consistent with the formation of a complex that remains stable in 0.5 M NaCl. The following strains were analyzed: T961.5h (*h4v-flag*) and T976.5h (*h4v-flag, asf1-ha*). **(B)** Ion-exchange chromatography (Q Sepharose) of lysates from strain T976.5h co-expressing H4v-FLAG and ASF1-HA. Proteins were eluted with an increasing NaCl gradient and analyzed by Western blotting as in Fig. 3A. A subset of ASF1-HA co-elutes with H4v-FLAG in fractions 5-6. **(C)** Gel filtration chromatography (Superdex 200) of whole-cell lysates from a *sad-6Δ* strain expressing H4v-FLAG (T965.4h). **(D)** Co-immunoprecipitation analysis of H4v-FLAG and ASF1-HA from Q Sepharose fractions 5 and 9. Interaction between H4v and ASF1 is readily detected in fraction 5 but not fraction 9. **(E)** Post-translational modifications on H4v identified by mass spectrometry. Acetylation sites corresponding to canonical H4K8 and H4K16 are shown together with variant-specific modifications, R7 methylation and K22 acetylation within the H4v-specific N-terminal extension. Tentative ubiquitination sites are also shown. The underlying protein, peptide, and PTM-level data are provided in the tables in Dataset 4.

Mass spectrometry of the H4v-containing complex immunoprecipitated from Superdex fraction 8 identified histone H3 and the chaperone ASF1 (Fig. S2D). To validate the interaction between H4v and ASF1, an HA epitope was introduced at the C-terminus of ASF1. ASF1-HA was found in Superdex fractions 8-10, roughly co-eluting with H4v-FLAG (Fig. 3A). To better understand the relationship between H4v and ASF1, we used ion-exchange chromatography on Q Sepharose as an orthogonal approach to gel filtration. Whole-cell lysates from a strain producing H4v-FLAG and ASF1-HA were applied to a HiTrap Q column, and elution was done with a 0.3-1.0 M NaCl gradient (Fig. 3B). H4v-FLAG and ASF1-HA eluted in fractions 5-6, accounting for a small portion of total ASF1-HA, while most ASF1-HA eluted in fractions 8-9 (Fig. 3B). H4v-FLAG was also present in fractions 9-15 (Fig. 3B). H4v-FLAG and ASF1-HA interacted in Q fraction 5 but not in fraction 9 (Fig. 3D). These results suggest that H4v, H3 and ASF1 form a complex, which represents a minor portion of the total ASF1-bound H3–H4 pool. Consistent with this idea, H3 and H4v were identified among the proteins associated with ASF1 in *Sordaria macrospora*^24^, and a direct interaction between H3 and H4v was reported in *Aspergillus nidulans*^22^.

We used gel filtration chromatography to test whether SAD-6 was required for the ASF1–H3–H4v complex. H4v-FLAG from a *sad-6Δ* strain continued to elute as a peak in Superdex fraction 8 (Fig. 3A,C), consistent with the ASF1–H3–H4v complex remaining in the absence of SAD-6. In parallel, an altered pattern of H4v-FLAG signal was noted in fractions 2–6 (Fig. 3A,C), suggesting that larger H4v-containing assemblies may become reorganized in the *sad-6Δ* condition, potentially contributing to the altered distribution of H4v-GFP nuclear signal observed by fluorescence microscopy (Fig. 2D).

### H4v carries conserved and variant-specific post-translational modifications

Mass spectrometry of post-translational modifications (PTMs) of H4v identified acetylation at K33 and K41, which correspond to canonical H4K8 and H4K16, respectively (Fig. 3E; Fig. S2A). Within the H4v-specific N-terminal extension, acetylation at K22 and methylation at R7 were also identified (Fig. 3E). Notably, R7 is found in an RGGP motif, a known target for protein arginine methyltransferases, which preferentially modify arginine residues in RG/RGG contexts^25^. Additionally, tentative ubiquitination was revealed at K45 and K120 (Fig. 3E). Thus, H4v carries both conserved and variant-specific PTMs, which may be involved in regulating its function.

### Several loci including *frequency* exhibit highly variable levels of H4v

While genome-wide H4v enrichment was largely congruent between the reference and *qde-1Δ* conditions, a few loci exhibited variable H4v profiles across six biological and technical replicates from both backgrounds (Fig. S3A,B). For some of these loci, such as NCU02421-NCU02423, NCU02664, NCU05568-NCU05570, NCU06853, NCU08594, and NCU10073, the variation appeared qualitative and lineage-specific rather than caused by the loss of QDE-1 (Fig. S3C). At other loci, including the promoter region of the gene *frequency* (*frq*/NCU02265), H4v levels varied more sporadically (Fig. S3C). Because *frq* is known to undergo dynamic H3K9me3 and 5mC linked to convergent transcription^12,13^, H4v enrichment at this locus pointed to a broader role for SAD-6 in chromatin regulation.

### H4v marks distinct types of accessible chromatin

H4v-enriched loci were identified with MACS3^26^ and divided initially according to their association with AT-rich DNA, which mainly corresponds to repetitive elements mutated by RIP. Peaks overlapping or bordering AT-rich sequences were classified as “AT-rich”, whereas the remaining peaks were classified as “GC-norm” (Fig. 4A; Fig. S4A). Of the ∼2,700 identified peaks, ∼1,700 were assigned to the GC-norm category. Overall, H4v peaks were distributed throughout the genome and were found at telomeres (Fig. S4B), rDNA loci (Fig. S4D), tRNA genes (Fig. S4C), as well as other genic and intergenic regions (Fig. 4D; Fig. S5).

**Figure 4.**
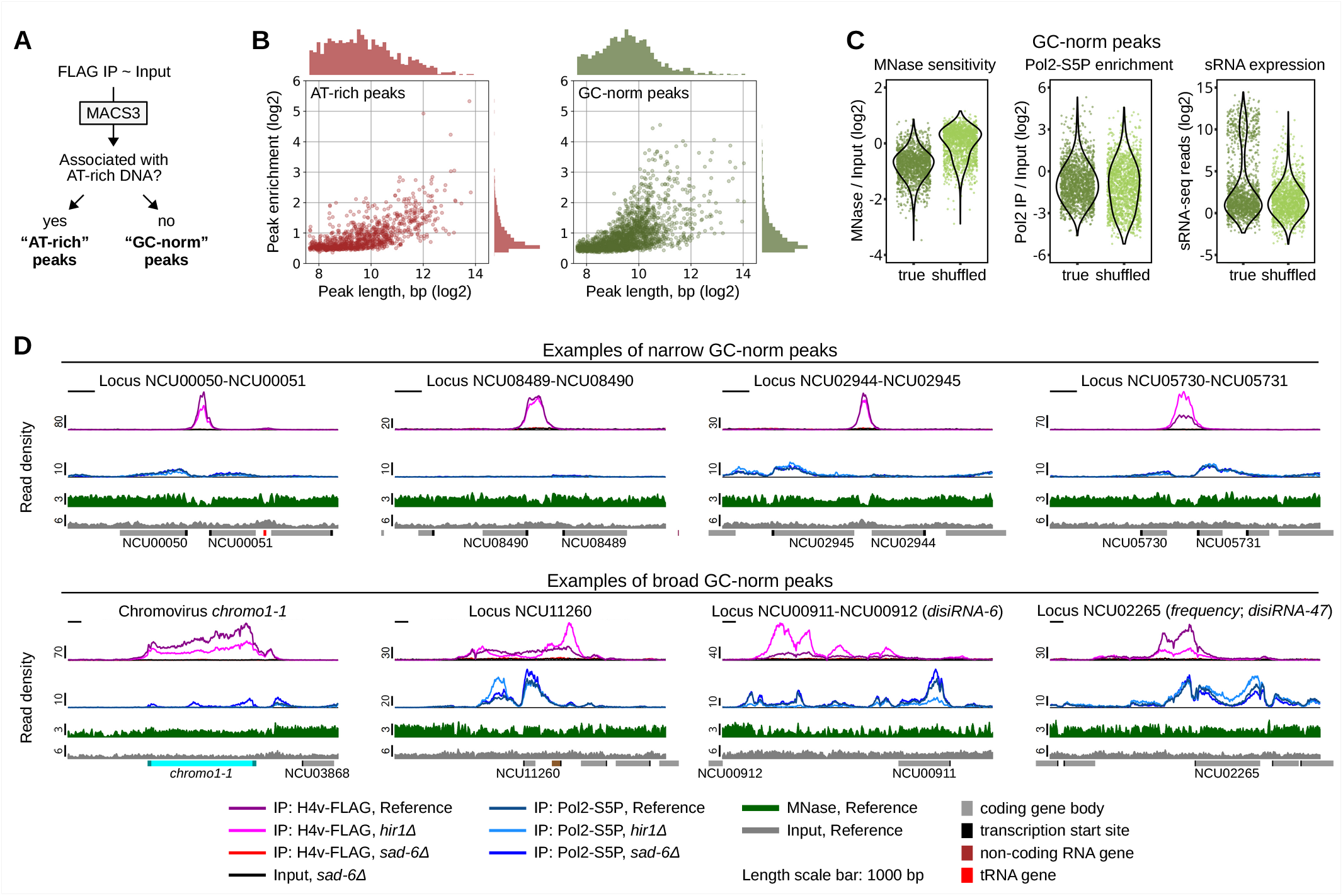
H4v is associated with distinct chromatin domains. **(A)** Schematic overview of the peak classification workflow. H4v-FLAG ChIP-seq peaks were identified using MACS3 and subsequently classified according to association with AT-rich DNA into “AT-rich” and “GC-norm” peak classes. The following strains were analyzed: T961.5h and T961.7h. **(B)** Relationship between peak length and enrichment for AT-rich and GC-norm peaks. Scatter plots show H4v enrichment as a function of peak length. Marginal histograms indicate the distributions of peak length and enrichment for each class. **(C)** Genomic and chromatin properties of GC-norm peaks, as compared to shuffled controls. Violin plots compare true GC-norm peaks and shuffled genomic intervals for MNase sensitivity (MNase/Input log2), Pol2-S5P enrichment (IP/Input log2), and small RNA abundance (sRNA-seq reads, log2). **(D)** Representative H4v-FLAG peaks. Tracks show H4v-FLAG ChIP-seq in several genetic backgrounds as indicated, together with corresponding input controls, Pol2-S5P ChIP-seq, and MNase-seq profiles. Gene annotations are shown below, including coding genes, transcription start sites, tRNA genes, and non-coding RNA genes. Examples are separated into narrow GC-norm peaks (top) and broad GC-norm peaks (bottom). Narrow peaks are typically highly localized and frequently associated with intergenic or promoter-proximal regions, whereas broad peaks span extended chromatin domains often associated with repetitive or atypical loci. Broad GC-norm peaks frequently coincide with structured MNase-seq patterns, suggesting association with distinctive chromatin organization. Analyzed strains are listed in Fig. 2E. Scale bar, 1000 bp.

We next investigated how H4v peaks differed in their profiles by comparing peak width versus enrichment across AT-rich and GC-norm categories (Fig. 4B). Peaks shorter than approximately 2 kbp (log2 length <11) were considered narrow, whereas larger peaks were considered broad. The vast majority of GC-norm peaks were narrow, while a higher proportion of AT-rich peaks had broad profiles. With respect to enrichment, GC-norm peaks showed somewhat higher values than AT-rich peaks; yet most peaks in both categories exhibited less than two-fold enrichment over input (Fig. 4B).

We focused on GC-norm peaks and assessed their association with specific chromatin features by comparing their distributions with those of randomly shuffled genomic controls (Fig. 4C). GC-norm peaks were strongly associated with MNase-sensitive regions yet, overall, did not show clear Pol2-S5P enrichment (Fig. 4C). The peaks were enriched for small RNAs; yet this pattern appeared linked to their overlap with tRNA genes (Fig. S4C), which represent a potent source of small RNAs in sRNA-seq libraries^9^.

Representative genome-browser views illustrate the distinct features of narrow versus broad GC-norm peaks (Fig. 4D). Narrow peaks display sharp, localized H4v enrichment and are typically found at short intergenic regions, with limited Pol2-S5P overlap. In contrast, broad peaks are frequently associated with complex patterns of MNase sensitivity and Pol2-S5P occupancy. Notably, this category includes *disiRNA* loci, including *disiRNA-6* and *disiRNA-47* (*frequency*), two well-characterized examples of dynamic H3K9me3 and 5mC^12^. Comparison of the reference and *hir1Δ* backgrounds showed that H4v localization was generally preserved in the absence of HIR1, whereas enrichment levels and peak shapes were often altered (Fig. 2F; Fig. 4D; Fig. S4B; Fig. S5), indicating that HIRA modulates H4v distribution (directly or indirectly) within the SAD-6-defined genomic intervals.

### SAD-6-dependent H3K9me3 marks a broad set of gene-proximal H4v loci

The association of H4v with known examples of dynamic H3K9me3 and 5mC encouraged us to evaluate this relationship in more detail. Inspection of prominent GC-norm peaks identified three classes of gene-proximal H4v loci based on their relationship with H3K9me3 (Fig. S5). The first class exhibited strong H3K9me3 that was SAD-6-independent (Fig. S5A), consistent with its induction by AT-rich sequence motifs^17^. Indeed, close inspection revealed local reductions in GC content (Fig. S5A), which nevertheless were above the threshold used for classification. The second class of H4v peaks displayed substantially lower levels of H3K9me3 that were dependent on SAD-6 (Fig. S5C). This group contained the model *disiRNA* loci, including *frq* (Fig. 5C). The third class comprised loci with robust H4v enrichment but no detectable H3K9me3 (Fig. S5B).

**Figure 5.**
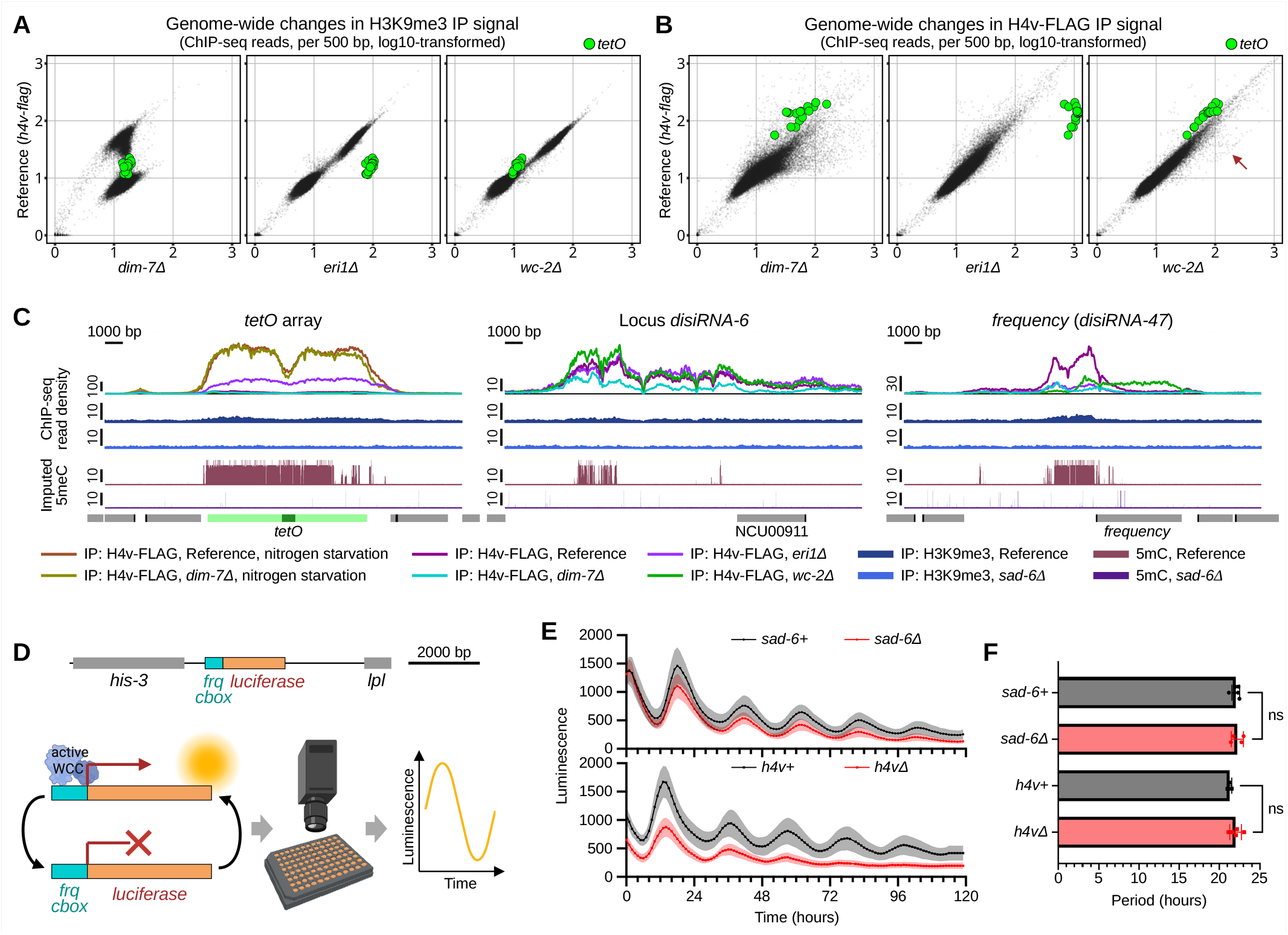
H4v deposition and associated H3K9me3 are influenced by diverse processes. **(A)** Genome-wide comparison of H3K9me3 ChIP-seq profiles between indicated genetic backgrounds, analyzed as in Fig. 1E. The following strains were analyzed: T961.5h and T961.7h (Reference); T971.2h and T971.3h (*dim-7Δ*); T974.6h and T974.7h (*eri1Δ*); T980.6h and T980.12h (*wc-2Δ*). **(B)** Genome-wide comparison of H4v-FLAG ChIP-seq profiles between indicated genetic backgrounds, analyzed as in Fig. 1E. Analyzed strains are listed in Fig. 5A. **(C)** ChIP-seq (H4v-FLAG and H3K9me3) and imputed 5mC profiles at *tetO* array, *disiRNA-6*, and *frequency* (*disiRNA-47*). Strains analyzed by ChIP-seq are listed in Fig. 5A; same *dim-7Δ* strains were also subjected to nitrogen starvation. The following strains were analyzed for 5mC: T961.5h (Reference); T965.4h (*sad-6Δ*). **(D)** Reporter system to assay core circadian clock activity. The C-box-containing region of the *frq* promoter was fused to the firefly luciferase gene, enabling monitoring of the WCC activity over several days using a 96-well plate format. **(E)** Luciferase assays of strains carrying *sad-6Δ* (top) or *h4vΔ* (bottom) and their corresponding wild-type controls. Each sibling strain represents one biological replicate and was assayed in 3-4 technical replicates. Per-sibling measurements were aggregated by genotype; solid lines indicate the mean and shaded regions represent one standard deviation. **(F)** Circadian periods estimated from the luciferase assays in Fig. 5E. Each point represents one sibling strain (biological replicate). Statistical significance was assessed by one-way ANOVA.

Because gene-proximal SAD-6-dependent H3K9me3 signal was generally weak, we sought to validate our observations using an orthogonal approach, by taking advantage of the fact that 5mC tracks H3K9me3 in *N. crassa*^17,27^. We therefore profiled 5mC from Oxford Nanopore long-read sequencing data. To avoid ambiguity associated with the dynamic nature of H3K9me3/5mC at the *frq* promoter^13^, great care was taken to process the strains in parallel. Consistent with the ChIP-seq results, deletion of *sad-6+* was accompanied by the loss of 5mC at the *tetO* array and at the model *disiRNA* loci (Fig. 5C).

### Genomic distribution of H4v is influenced by H3K9me3

Our results revealed several types of loci at which H3K9me3 enrichment depends on the SAD-6/H4v system. Thus, we next tested whether H3K9me3, in turn, affects H4v enrichment. We used deletion of *dim-7+*, which encodes a key component of the H3K9me3 pathway in *N. crassa* (DIM-7)^17^, to eliminate H3K9me3 genome-wide (Fig. 5A). Remarkably, deletion of *dim-7+* coincided with a large-scale redistribution of the H4v signal (Fig. 5B). For example, H4v enrichment was reduced at the unperturbed *tetO* array, model *disiRNA* loci, and some genes (NCU06853 and NCU11260), and it was nearly eliminated from several short intergenic regions that normally exhibited H3K9me3, including the intervals NCU01005-NCU01006, NCU03305-NCU03306, and NCU06149-NCU06150 (Fig. 5B,C; Fig. S6A). At the same time, many prominent H4v peaks emerged in AT-rich regions, either within AT-rich blocks or at the boundaries between AT-rich and AT-normal DNA (Fig. S6A). We also investigated whether H3K9me3 was required for the remarkably strong H4v enrichment at the perturbed *tetO* array and found that loss of DIM-7 did not impair H4v accumulation following acute nitrogen starvation (Fig. 5C).

Together, our results reveal a context-dependent relationship between H3K9me3 and H4v, with H3K9me3 ranging from dispensable to essential for H4v enrichment and exerting both positive and negative effects. More generally, although H4v deposition absolutely requires SAD-6, its genomic distribution is ultimately shaped by additional chromatin processes.

### ERI1 modulates H4v enrichment in a locus-dependent manner

The RNA exonuclease ERI1 was identified as a positive regulator of H3K9me3 and 5mC at *frq*^28^. Consistent with this role, loss of ERI1 resulted in a strong reduction of H4v enrichment at *frq* (Fig. 5C). However, this requirement was not universal, as *disiRNA-6*, the other model *disiRNA* locus, still exhibited wild-type levels of H4v in the *eri1Δ* condition (Fig. 5C). Remarkably, loss of ERI1 provoked a dramatic increase in H4v and H3K9me3 enrichment at the *tetO* array, assayed in the absence of TetR-GFP or nitrogen starvation (Fig. 5A-C). The effect was highly specific to the *tetO* locus, as comparable gains were not observed elsewhere in the genome (Fig. 5A-B). This result is consistent with previous work reporting elevated siRNA production from the repressor-bound *tetO* array in the absence of ERI1 in *N. crassa*^9^, and it also parallels the role of ERI1 as a negative regulator of exogenous RNAi in *C. elegans*^29^. Together, these findings indicate that ERI1 modulates SAD-6-dependent H4v-associated chromatin in a locus-specific manner. Whether these effects reflect direct regulation of SAD-6 activity or indirect consequences of altered RNA metabolism remains to be determined.

### WC-2 is required for H4v enrichment at the *frq* promoter

Periodic transcription of the *frq* gene is driven by the White Collar Complex (WCC) composed of WC-1 and WC-2^30^. WC-2 binds the Clock box (C-box) within the *frq* promoter through its GATA zinc-finger domain^30^. Previous studies found that WC-2 was required for 5mC at the *frq* promoter, suggesting that WCC-dependent transcription promotes a chromatin environment permissive for dynamic H3K9me3 deposition^12^. Our results show that the formation of this dynamic H3K9me3 requires SAD-6 (Fig. 5C). These observations raised the possibility that H4v enrichment at the *frq* promoter also depends on WC-2. Indeed, we found that deletion of *wc-2+* strongly reduced H4v signal across this interval (Fig. 5C). This effect was specific, as the other model *disiRNA* locus (*disiRNA-6*) retained normal levels of H4v in the *wc-2Δ* condition (Fig. 5C).

Unexpectedly, loss of promoter-associated H4v was accompanied by the emergence of a new H4v peak over the *frq* coding region (Fig. 5C). This peak was present in only one of the two *wc-2Δ* strains used as biological replicates (Fig. S6B). Genome-wide analysis identified additional intervals that gained H4v in the absence of WC-2 (Fig. 5B). Inspection of the corresponding ChIP-seq profiles for a subset of the affected loci revealed that these gains occurred sporadically in either one or the other *wc-2Δ* strain (Fig. S6B), suggesting that loss of WC-2 was associated with stochastic redistribution of H4v at a limited number of sites. Notably, one such site corresponds to the DNA transposon *sly1-1*^31^, a target of the clock-controlled transcription factor ADV-1^32^, while another site contains a gene encoding the chromatin remodeler CHD1 (NCU03060), a known regulator of dynamic H3K9me3 at *frq*^13^. While these patterns are intriguing, they may reflect the inherently stochastic nature of H4v enrichment at some genomic intervals (Fig. S3), rather than WC-2 loss.

### Circadian clock operates normally in the absence of SAD-6 and H4v

Because H4v, H3K9me3 and 5mC at the *frq* promoter required SAD-6, we next investigated whether SAD-6 and H4v are also required for clock function. We used a previously developed reporter containing the C-box DNA element from the *frq* promoter fused to the firefly *luciferase* gene (*frq_cbox_-luc*), which allowed tracking clock activity in real time and over a period of several days^33^ (Fig. 5D). For each condition (*sad-6Δ* or *h4vΔ*), a panel of several reporter strains was established by crossing a corresponding deletion strain to the *frq_cbox_-luc* strain 661-4A^33^. Overall, the panels included six *sad-6+* and four *sad-6Δ* siblings, as well as three *h4v+* and five *h4vΔ* siblings. For each sibling, luminescence curves and circadian period estimates were based on 3-4 technical replicates assayed in the 96-well format. Per-sibling measurements were aggregated by genotype for statistical analysis and visualization.

Luminescence recordings revealed robust circadian oscillations in all genotypes (Fig. 5E,F). Neither *sad-6Δ* nor *h4vΔ* significantly altered circadian period or overall reporter cycling, suggesting that loss of either factor does not measurably affect WCC activity. These results indicate that dynamic H3K9me3 and 5mC at *frq* arise from a SAD-6-dependent response to WCC-driven transcription and are dispensable for core clock function, consistent with a model in which these marks represent a downstream consequence of transcription-coupled chromatin remodeling rather than feedback regulation of the circadian clock.

### SAD-6 targets foreign DNA that escaped effective RIP

At the perturbed *tetO* array, SAD-6-dependent H3K9me3 reached levels comparable to those observed in AT-rich constitutive heterochromatin. In contrast, at most other loci, SAD-6-dependent H3K9me3 was present at much lower levels, consistent with its dynamic nature. Previously, we described a region (*r2*) in which a GC-normal sequence embedded in an AT-rich block featured robust SAD-6-dependent H3K9me3, comparable to that at the perturbed *tetO* array and in constitutive heterochromatin^9^. We have now determined that this locus corresponds to a full-length copy of a chromovirus, which we therefore named *chromo1-1* (Fig. 4D). Overall, *chromo1-1* exhibits strong H4v enrichment (Fig. S7A), which is regulated by additional processes, including the H3K9me3 pathway (Fig. 4D; Fig. S7B). Sequence analysis identified several point mutations distributed across the entire element, consistent with a history of mild RIP. For example, its two 289-bp LTRs contained three C-to-T transitions and no other mutation types. Yet *chromo1-1* retains an overall GC content of 50.5%, suggesting that its strong association with H4v and H3K9me3 is not simply a consequence of being AT-rich.

Deletion of *sad-6+* or *h4v+* resulted in a similar pattern of H3K9me3 loss over the element, accompanied by increased Pol2-S5P occupancy in its central part and near the LTRs (Fig. S7B). Interestingly, Pol2-S5P levels increased at the LTR that remained associated with H3K9me3, indicating that H3K9me3 alone is insufficient to maintain robust transcriptional silencing of *chromo1-1* without the SAD-6/H4v system.

Because *chromo1-1* is found in the constitutive H3K9me3 block, it remained unclear whether its exceptional enrichment in H4v and H3K9me3 was simply a consequence of its genomic position. To test this possibility, we replaced its “left” LTR with a 6247-bp sequence containing the *ntcR* transcription unit oriented toward the AT-rich DNA block and flanked by two 2500-bp inserts of DNA from *E. coli* (Fig. S7C). Three effects were observed. First, the modified element featured unexpectedly high levels of H4v and H3K9me3. Second, H4v and H3K9me3 enrichment was even higher over the segment of *E. coli* DNA separating *chromo1-1* and *ntcR*. Third, a prominent H4v peak was formed at the boundary between AT-rich and *E. coli* DNA, coincident with Pol2-S5P enrichment (Fig. S7C). Taken together, these results indicate that SAD-6 can target *chromo1-1* for robust silencing even within heterochromatin, and such silencing can extend into nearby foreign DNA. The formation of the H4v peak at the AT-rich/*E. coli* DNA boundary suggests that transcription may contribute to H4v recruitment at chromatin transition zones.

### SAD-6 is absolutely required for DIM-2-dependent RIP

RIP is a process in which gene-sized DNA repeats trigger extensive C-to-T mutation in both themselves and adjacent genomic regions^15^. Studies in *N. crassa* suggest that RIP recognizes repeats via homologous pairing of intact DNA double helices^15^. RIP mutation is mediated by two largely independent pathways with distinct substrate preferences: RID-dependent RIP targets the repeats, whereas DIM-2-dependent RIP mostly affects the adjacent regions^15^ (Fig. 6A). DIM-2-dependent RIP also requires all major components of the canonical constitutive heterochromatin pathway^17,18^, effectively transforming transient repeat-induced H3K9me3 into a permanent mutational record. We therefore asked whether SAD-6 was required for DIM-2-dependent RIP.

**Figure 6.**
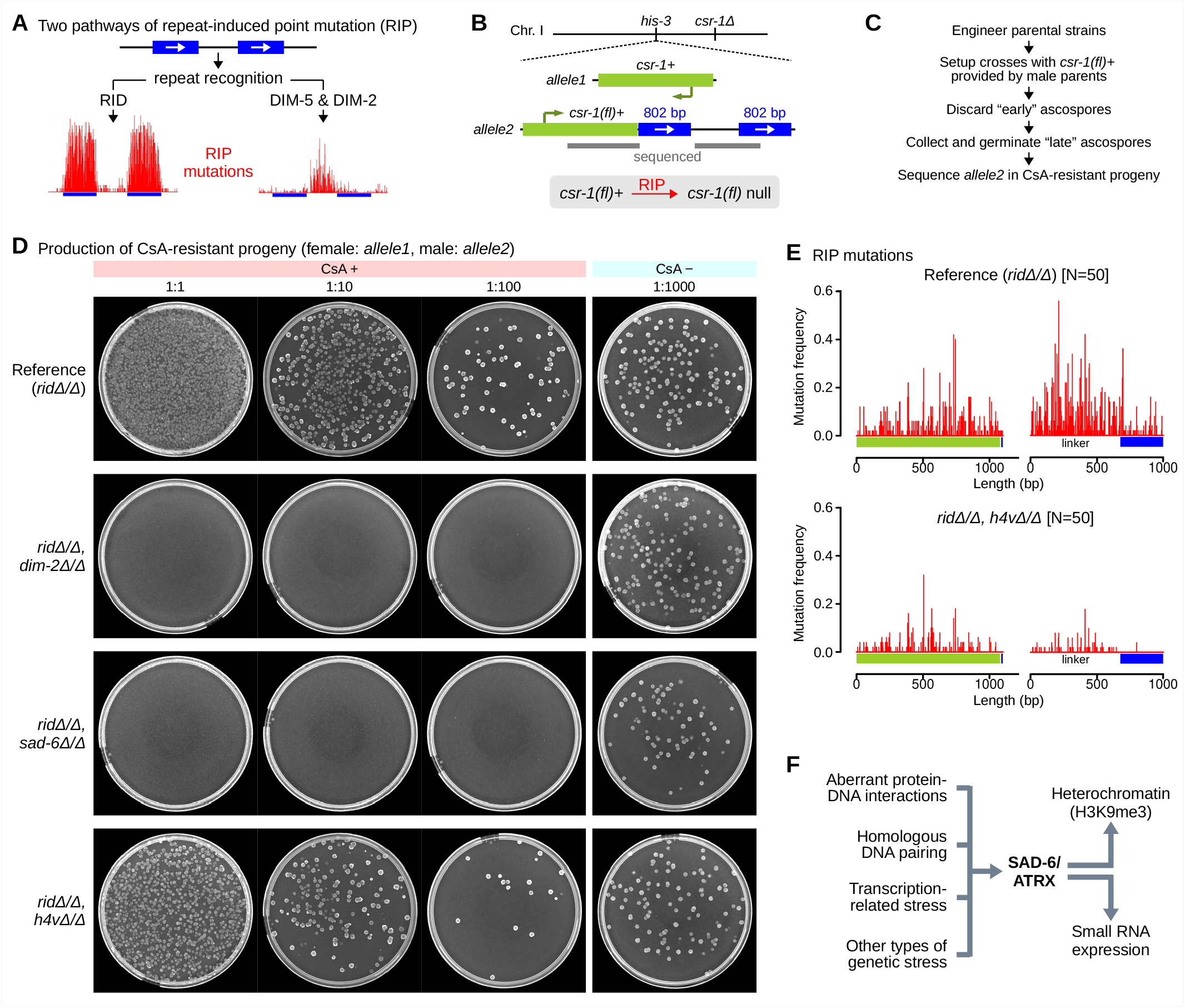
SAD-6 is required for DIM-2-dependent repeat-induced point mutation (RIP). **(A)** Overview of the two RIP pathways in *N. crassa*. Whereas RID-dependent RIP is concentrated in DNA repeats, DIM-5(SUV39)/DIM-2-dependent RIP preferentially mutates repeat-proximal regions. **(B)** Genetic system to detect DIM-2-dependent RIP. Mutation of the repeat-proximal *csr-1+* (*allele2*) yields cyclosporin A (CsA)-resistant progeny that can be readily recovered on selective medium. Gray bars denote regions analyzed by sequencing in Fig. 6E. **(C)** Experimental strategy for assaying DIM-2-dependent RIP using the system in Fig. 6B. **(D)** Production of CsA-resistant progeny in the indicated genetic backgrounds. Serial dilutions of ascospores were plated on selective and non-selective medium as indicated. Crosses between the following strains were assayed (listed as female x male parents): T804.8h x T805.1h (Reference); T998.4h x T1002.2h and T998.6h x T1002.5h (*dim-2Δ/Δ*); T996.5h x T1000.2h and T996.8h x T1000.12h (*sad-6Δ/Δ*); T997.7h x T1001.3h and T997.8h x T1001.4h (*h4vΔ/Δ*). Representative results are shown. Analyzed crosses are listed in Table S5. **(E)** Distribution of RIP mutations across the reporter locus in the indicated genetic backgrounds. Mutation frequency is plotted as a function of position along the repeat construct. Analyzed multiple sequence alignments are provided in Dataset 5, referenced by cross names listed in Table S5. **(F)** Model for SAD-6/ATRX function in fungal genome surveillance and defense. SAD-6 connects several types of genetic stress to heterochromatin formation and small RNA production, enabling efficient genome surveillance during vegetative growth and DIM-2-dependent RIP and MSUD during sexual reproduction.

We first developed a sensitive genetic system to assay DIM-2-dependent RIP (Fig. 6B). In this system, a pair of direct GC-rich repeats of *E. coli* DNA acted as a strong inducer of RIP^34^, whereas mutation of the adjacent *csr-1+* gene provided a sensitive readout, as it confers resistance to cyclosporin A in the progeny. Resistance was scored by plating germinated ascospores on selective medium, and mutations were confirmed by Sanger sequencing^18^. The system was implemented in a pair of strains carrying deletions of the endogenous *rid+* and *csr-1+* genes. The first strain (used as a female parent) also carried an ectopic copy of *csr-1+* near *his-3* (Fig. 6B: *allele1*). In the second strain, this position was occupied by the reporter construct (Fig. 6B: *allele2*). RIP was scored by plating original suspension and two ten-fold dilutions of ascospores on selective medium, with the final dilution plated on non-selective medium as a control (Fig. 6C,D).

The system was validated by testing its dependence on DIM-2. A reference *dim-2+/+* cross yielded abundant cyclosporin-resistant progeny (Fig. 6D). Sequence analysis of the reporter locus in those progeny identified a large number of C-to-T (and G-to-A) transitions (Fig. 6E). Other mutations were not detected. In contrast, no cyclosporin-resistant progeny were produced in the corresponding *dim-2Δ/Δ* condition (Fig. 6D). Increasing the number of analyzed ascospores to approximately one million still failed to recover any cyclosporin-resistant progeny, underscoring the sensitivity and specificity of this assay as a readout of DIM-2-dependent RIP.

Critically, loss of SAD-6 eliminated all RIP in this system, effectively recapitulating the *dim-2Δ/Δ* condition (Fig. 6D). RIP still occurred in the absence of H4v, yet it was markedly reduced (Fig. 6D,E). This defect was less pronounced in the flank, likely because our selection strategy relies on mutations to occur in this region.

To test whether SAD-6 is required for RIP in general, we modified the reporter construct by replacing the linker and the distal GC-rich repeat with a 950-bp segment of *csr-1*, thereby redefining *csr-1+* as an inverted repeat and making it a better substrate for RID-dependent RIP (Fig. S8A; compare *allele2* and *allele3*). This modification also redefined the remaining GC-rich repeat as a linker (Fig. S8A). Curiously, the *sad-6Δ* strain carrying this inverted repeat appeared resistant to cyclosporin before sexual crossing and associated RIP (Fig. S8B). The irregular colony morphology pointed to an epigenetic basis for this phenotype. Indeed, abundant siRNAs (originating mostly from the duplication but also from the linker) were detected, implicating RNAi as the cause of resistance (Fig. S8C-E). A cross lacking RID and SAD-6 yielded many cyclosporin-resistant progeny (Fig. S8F). Seventy such progeny were chosen for analysis, and all were found to contain the repeat construct, suggesting that resistance remained linked to the silenced *csr-1* allele after meiosis. Sequencing of the *csr-1* duplication in this sample showed no evidence of RIP (Fig. S8G: *ridΔ/Δ, sad-6Δ/Δ*). Crucially, RIP was restored by replacing *ridΔ* with *rid+* in one parental strain (Fig. S8F,G: *rid+/Δ, sad-6Δ/Δ*). These results indicate that SAD-6 controls the heterochromatin-related (DIM-2-dependent) pathway of RIP, whereas repeat recognition and RID-dependent RIP can still occur in its absence.

## DISCUSSION

### ATRX-like chromatin remodelers may not universally require H3.3

The role of ATRX in promoting gene silencing has been linked to replication-independent deposition of H3.3 at dynamic chromatin regions, including repetitive DNA and loci associated with non-B-DNA structures^2^. In animals, this activity is mediated by the ATRX–DAXX complex^2^. However, the evolutionary distribution of these factors suggests that this mechanism cannot be universal: while ATRX orthologs are broadly conserved in eukaryotes, DAXX is not conserved in plants^7^ and fungi^10^, and dedicated H3.3 variants are also lacking in many fungal species^10,23^. Our results resolve this apparent paradox by showing that the fungal ATRX ortholog SAD-6 instead relies on the histone H4 variant H4v for its function in genome surveillance and defense.

Our results indicate that the histone chaperone requirements of ATRX-like remodelers can be more flexible than previously thought. We find that H4v forms a complex with ASF1 (Fig. 3), a highly conserved histone chaperone that serves as a hub for transferring H3–H4 dimers to several downstream chaperones, including DAXX, HIRA, and CAF-1^35^. The lack of requirement for HIRA (Fig. 2), together with the apparent absence of DAXX in fungi, suggests that SAD-6-mediated H4v deposition relies on an ASF1-dependent pathway that does not involve canonical H3.3-specific chaperones. This raises the possibility that ASF1-associated histone complexes are either directly engaged by SAD-6 or transferred to SAD-6 via an additional factor. Consistent with this model, the absolute requirement of DIM-2-dependent RIP for SAD-6 contrasts with its only partial requirement for H4v, suggesting that SAD-6 may potentially utilize histone chaperone complexes containing canonical H3–H4 dimers.

### H4v is a highly divergent H4 variant involved in genome surveillance and defense

Our identification of H4v as a cofactor of SAD-6 expands the functional repertoire of histone H4 variants. H4v was first described in *N. crassa*^10,21^ and later found to be broadly conserved across the Pezizomycotina^22^. Other H4 variants include H4V in trypanosomes, which marks transcription termination sites of polycistronic transcription units^36^; H4G in humans and some other primates, which localizes to nucleoli and functions as a chromatin-associated factor rather than a replacement for H4^37^; and H4.V in rice, implicated in modulation of chromatin in response to developmental or environmental cues^38^. These variants share ∼73-85% identity with their canonical counterparts and lack a C-terminal extension predicted to form a novel histone domain. Thus, H4v is distinguished among H4 variants by its remarkable structural divergence, conservation over hundreds of millions of years of evolution, and its pervasive role in genome surveillance and defense.

### SAD-6 enables broad genome surveillance

SAD-6 was first described in *N. crassa* as a factor required for robust MSUD^8^ and was later found to control H3K9me3 deposition and small RNA production in response to experimental perturbation of chromatin^9^. Our findings expand the functional repertoire of SAD-6 and suggest that fungal ATRX-like chromatin remodelers play broad genome regulatory roles. Genome-wide profiling identified nearly 2,700 peaks of H4v associated with both coding and non-coding regions. H4v enrichment at all identified peaks required SAD-6. The most apparent feature of these peaks was their MNase sensitivity, suggesting that they mark regions of structurally dynamic chromatin. The peaks could be grouped by their size and genomic context. Narrow peaks (<∼2 kbp) were more abundant than broad peaks (>∼2 kbp), being situated between closely-positioned genes (irrespective of gene orientation), tRNA loci, and also the boundaries between AT-rich and AT-normal domains. Broad peaks typically spanned extended intergenic regions but were also occasionally observed across large genes.

In contrast to the *tetO* array, which may potentially exist in the perturbed state in the majority of nuclei, most H4v peaks feature much lower enrichment, indicating that they are occupied by H4v-containing nucleosomes in only a subset of nuclei. Consistent with this model, several peaks displayed substantial variability between biological and technical replicates, indicative of a highly dynamic, and potentially stochastic, pattern of H4v deposition. Notably, the overall genome-wide distribution of H4v in *N. crassa* resembles that of mammalian ATRX (including enrichment at subsets of coding and promoter-proximal regions^3^), suggesting that the broad regulatory roles of ATRX-like remodelers may have been conserved across fungi and animals. Yet the lack of clear developmental defects in *N. crassa* strains deficient in SAD-6/H4v indicates that this system evolved to serve a more specialized role in genome surveillance and defense in fungi (Fig. 6F).

In this study, H4v enrichment served as a genome-wide readout of SAD-6 activity, revealing its operation at telomeres, tRNA and rDNA loci, decaying mobile elements that escaped hypermutation by RIP, AT-rich DNA domains, and many gene-proximal intervals. Gene-proximal H4v peaks could be classified according to their relationship with H3K9me3. The first class corresponded to H4v peaks featuring strong, SAD-6-independent H3K9me3, consistent with its deposition by a canonical mechanism linked to AT-rich sequence motifs^17^. The second class was associated with much weaker, SAD-6-dependent H3K9me3; whereas the third class lacked detectable H3K9me3. The division between the second and third classes was likely not absolute, reflecting, at least in part, the sensitivity limits of H3K9me3 detection by ChIP-seq. The second class included several loci previously shown to generate small RNAs and undergo dynamic H3K9me3 and 5mC, including the *frq* gene that controls the circadian clock in *N. crassa*^11–13^. Yet loss of SAD-6 and H4v did not affect core clock function, suggesting that the dynamic H3K9me3 and 5mC states at the *frq* locus may represent a downstream outcome of transcription-induced chromatin remodeling rather than essential feedback regulators.

### SAD-6 links repeat recognition via homologous DNA–DNA pairing to heterochromatin nucleation

The association of repetitive DNA with H3K9me3-rich heterochromatin is a landmark feature of nearly all eukaryotic genomes^39,40^. In animals, formation of repeat-associated heterochromatin is RNAi-independent and requires, among other factors, the SUV39H1/2 lysine methyltransferases and ATRX^2,39^. How repetitive DNA is recognized for silencing in the absence of strong sequence-specific signals remains unclear. Current models invoke the formation of various nucleic acid structures, including R-loops and cruciforms, which are enriched at repetitive sequences and have been linked to genome instability and chromatin regulation^39,41,42^. Another proposal involves higher-order DNA complexes produced by recombination-independent pairing of homologous DNA double helices, which could trigger sequence-independent heterochromatin nucleation on repetitive DNA^18,43^.

The strongest evidence for DNA homology recognition through recombination-independent pairing comes from genetic studies of RIP and MSUD in *N. crassa*^15^. Both processes can detect homology with remarkable specificity and efficiency while operating independently of RecA-like recombinases^43,44^. Analysis of DIM-2-dependent RIP at closely-positioned repeats showed that direct repeats induce substantially stronger mutation in adjacent regions than inverted repeats, consistent with propagation of a local topological response^34^. These observations have been interpreted as supporting a model in which homologous DNA pairing generates local supercoiling stress that is dissipated more efficiently in inverted than in direct repeat configurations, thereby enhancing DIM-2-dependent RIP near direct repeats. Our finding that DIM-2-dependent RIP requires SAD-6 suggests that ATRX-like chromatin remodelers may similarly promote heterochromatin formation by acting downstream of homologous DNA pairing, for example by responding to pairing-induced topological stress. Together with the requirement for SAD-6 in efficient MSUD^8^, this result supports a broader role for ATRX-like remodelers in coupling homologous DNA–DNA interactions to diverse downstream genome regulatory pathways.

## ACKNOWLEDGMENTS

This work was supported by the Agence nationale de la recherche (ANR-10-LABX-0062, ANR-11-LABX-0044, ANR-10-IDEX-0001-02, ANR-19-CE12-0002), the National Institutes of Health (R35GM118021), the Research Council of Finland (#321584) and Institut Pasteur. We acknowledge the Cell and Tissue Imaging Platform “PICT-IBiSA” (funded by ANR-10-INBS-04) and technical assistance of Jeanne Brouillet, Jeanne Contal, Sebastian Castro Ramirez, and Jasmin Ostermayer.

## AUTHOR CONTRIBUTIONS

F.C. and E.G. designed the study; F.C., A.K., E.B., Z.W., I.L., A.T., I.K., J.C.D., M.M., and E.G. performed experiments and analyzed the data; E.G. wrote the manuscript. All the authors have read and approved the manuscript.

## DECLARATION OF INTERESTS

The authors declare no competing interests.

## DATA AVAILABILITY

Sequencing data generated in this study were submitted to the Sequence Read Archive (SRA) under accession PRJNA1492767. Additional datasets are available at Figshare (DOI: 10.6084/m9.figshare.32952437).

## MATERIALS AND METHODS

### Manipulation of *Neurospora* strains

#### A. Strains

*N. crassa* genes analyzed in this study are listed in Table S1. *N. crassa* strains used in this study are listed in Table S2.

#### B. Growth media

VM medium (1× Vogel’s Medium N salts, 1.5% sucrose) was used for standard vegetative growth. Sorbose medium (1× VM salts, 2% sorbose, 0.1% dextrose, 3% agar) was used for colonial growth. If needed, media were supplemented with 5 µg/mL cyclosporin A (Sigma-Aldrich, cat. no. 30024; to select for *csr-1* progeny), 100 µg/mL tetracycline (Sigma-Aldrich, cat. no. T7660; to dissociate TetR-GFP from the *tetO* array), or 0.1 µg/mL blasticidin S (MP Biomedicals, cat. no. 150477; to induce expression of FLAG-SAD-6). For nitrogen starvation, VM medium was prepared without ammonium nitrate and used for the last 5 h of liquid culture as previously described^9^.

#### C. Transformations

Macroconidia (20 µL, washed and pelleted in 1 M sorbitol) were combined with 2-3 µg of linearized plasmid DNA in 10 µL of 1 M sorbitol, incubated on ice for 20 min, and electroporated using the following settings: 1500 V, 600 fi, 25 µF, 2 mm gap. Immediately after electroporation, macroconidia were resuspended in 1 mL of 1 M sorbitol. For transformations with plasmids targeting *his-3* or *csr-1*, macroconidia were plated directly onto sorbose agar. For transformations with plasmids carrying resistance markers for G418 or nourseothricin, macroconidia were allowed to recover for 3-4 h at ambient temperature before being plated onto sorbose agar containing 100 µg/mL G418 sulfate (Euromedex, cat. no. EU0601) or 50 µg/mL nourseothricin (Jena Bioscience, cat. no. AB-101L). Colonies were picked using glass Pasteur pipettes. Genomic DNA was extracted as previously described^43^, and integration events were analyzed by PCR. Homokaryotic strains were obtained from primary transformants by macroconidiation. Complete sequences of all transformation plasmids used in this study are provided in Dataset 2.

#### D. Linear vegetative growth assay

Macroconidia (∼10⁴ CFUs in 1 µL of water) were spotted at the center of 90-mm Petri plates (Gosselin, cat. no. SB90-101) containing VM medium solidified with 1.5% agar. If required, nourseothricin was used at a concentration of 50 µg/mL. Inoculations were performed between 17:30 and 18:00. Plates were incubated overnight at 32 °C, and initial growth was recorded the following morning (9:00-10:00) by tracing the edge of the mycelium on the underside of the plate. Plates were then returned to 32 °C for continued incubation. A second measurement was taken 10 hours later using the same procedure, after which the agar medium was discarded. Three lines passing through the inoculation point were drawn at random angles on the underside of each plate, generating six radial segments between the two recorded colony contours. Linear growth was calculated as the mean length of these six segments divided by 10 hours.

### Selection of nourseothricin-resistant mutants

#### A. Generation of clonal populations of macroconidia

Microconidia of the reference strain T838.4 were produced on a nitrogen-poor medium supplemented with 1 mM sodium iodoacetate^45^. Microconidia were purified by filtration through a 5-µm syringe filter (Merck, cat. no. SLSV025LS) and plated at low dilution on sorbose agar. Colonies were picked with Pasteur pipettes and used to initiate clonal cultures in glass tubes (17 × 180 mm), containing 3 mL of VM medium solidified with 1.5% agar. Cultures were kept at 24 °C for 10 days to allow macroconidia production. Mature macroconidia (4-6 × 10⁸ CFUs) were harvested in water, divided into eight aliquots, pelleted, and stored at −80 °C.

#### B. Selection of primary nourseothricin-resistant isolates

Each aliquot of macroconidia was spread on 12 90-mm Petri plates (Gosselin, cat. no. SB90-101) containing sorbose agar supplemented with 50 µg/mL nourseothricin. After drying, plates were UV-irradiated at 2 mJ/ cmZ using a GS Gene Linker (Bio-Rad) and kept in the dark at 24°C for 2 h to prevent photolyase-mediated DNA repair. Plates were subsequently incubated at 32°C for 2-3 days, after which colonies were picked with Pasteur pipettes and used to initiate primary cultures of nourseothricin-resistant isolates.

#### C. Phenotypic screening

Each primary isolate was assayed for linear growth in the absence and presence of nourseothricin. Isolates exhibiting growth rates above 0.5 mm/h in the presence of nourseothricin were selected for further analysis. Four homokaryotic lineages were obtained from selected primary isolates via microconidiation. Two of these homokaryotic isolates were re-assayed for linear growth under both selective and non-selective conditions. A subset of promising candidates was selected for Illumina sequencing. Two homokaryotic isolates per primary isolate were analyzed.

### Preparation of genomic DNA libraries for Illumina sequencing

Genomic DNA was extracted from small-scale mycelial cultures grown overnight in 2 mL of VM medium, as previously described^43^. Samples were treated with RNase A (Thermo Fisher Scientific, cat. no. EN0531), extracted once with phenol-chloroform, precipitated with 0.6 volumes of isopropanol, washed with 75% ethanol, and resuspended in TE buffer. DNA quality was assessed on agarose gels. DNA concentration was measured on a Qubit using the dsDNA HS Assay kit (Invitrogen, cat. no. Q33230) and adjusted in TE buffer to 10 ng/µL in a final volume of 500 µL. Samples were sonicated on ice using a Vibracell 75043 instrument (Bioblock Scientific) with the following program: total time = 2 min, pulse on = 2 s, pulse off = 8 s, 20% amplitude. Fragmented DNA (30 µL) was purified using Agencourt AMPure XP beads (Beckman Coulter, cat. no. A63880) according to the manufacturer’s instructions and eluted in 30 µL. Libraries were generated from 1-2 ng of eluted DNA using the NEBNext Ultra II DNA Library Prep Kit (NEB, cat. no. E7645S) with 11 cycles of PCR amplification.

### Chromatin immunoprecipitation (ChIP)

Large-scale cultures were initiated by inoculating approximately 1 × 10⁶ thawed macroconidia into 200 mL of VM medium. Cultures were grown for 24 h at 30 °C with shaking at 160 rpm. Mycelia were collected by filtration, washed with phosphate-buffered saline (PBS) at ambient temperature, blotted dry, and transferred to 100 mL of PBS containing 1% formaldehyde (Sigma-Aldrich, cat. no. 252549). Crosslinking was carried out for 30 min at ambient temperature with constant stirring and quenched by addition of glycine for 5 min. Mycelia were then collected by filtration, rinsed with PBS, blotted dry, frozen in liquid nitrogen, ground to a fine powder, and stored at −80 °C. The same mycelial preparation was used for both ChIP-seq and MNase-seq experiments.

Crosslinked mycelial powder (450 mg) was combined with 3 mL of sonication buffer (50 mM HEPES, pH 7.5, 140 mM NaCl, 1 mM EDTA, 1% Triton X-100, 0.1% sodium deoxycholate, and protease inhibitors [20 µL/mL, Roche, cat. no. 11873580001]) and sonicated on ice using a Vibracell 75043 instrument (Bioblock Scientific) with the following program: total time = 2 min, pulse on = 2 s, pulse off = 8 s, 40% amplitude. Lysates were cleared by centrifugation, twice at 13,000 × g for 5 min at 4°C. A 20-µL aliquot was retained as “Input” control.

For immunoprecipitation, cleared lysates were incubated overnight at 4°C with antibodies (1 µL of antibody per 1 mL of lysate) on a rotating wheel. Magnetic beads (Invitrogen, cat. no. 10746713) were washed twice in sonication buffer and added to the samples (20 µL beads per 1 mL of lysate); followed by incubation for 4 h at 4°C on a rotating wheel. Beads were sequentially washed (1 mL of buffer per wash, 10 min at 4°C) twice with lysis buffer (50 mM HEPES, pH 7.5, 140 mM NaCl, 1 mM EDTA, 1% Triton X-100, 0.1% sodium deoxycholate), once with high-salt buffer (50 mM HEPES, pH 7.5, 500 mM NaCl, 1 mM EDTA, 1% Triton X-100, 0.1% sodium deoxycholate), once with LiCl buffer (10 mM Tris-HCl, pH 8.0, 250 mM LiCl, 1 mM EDTA, 0.5% IGEPAL, 0.5% sodium deoxycholate), and once with TE buffer (10 mM Tris-HCl, pH 7.5, 1 mM EDTA). Antibodies used in this study are listed in Table S3.

Chromatin was eluted twice with 62.5 µL of TES buffer (50 mM Tris-HCl, pH 8.0, 10 mM EDTA, 1% SDS) at 65°C for 10 min, and eluates were pooled (125 µL total). Input samples were adjusted to the same volume with TES buffer. Crosslink reversal was done overnight at 65°C. Samples were treated with RNase A (2.5 µL, 2 h at 50°C), followed by proteinase K (6.25 µL of 20 mg/mL, 2 h at 50°C).

DNA was purified by phenol-chloroform extraction and ethanol precipitation (22 µL of 3 M sodium acetate, and 610 µL ethanol), incubated overnight at −20°C, precipitated by centrifugation at 20,000 × g for 30 min at 4°C, washed with 75% ethanol, air-dried, and resuspended in 20 µL of TE buffer. DNA concentration was measured on a Qubit using the dsDNA HS Assay kit (Invitrogen, cat. no. Q33230). Libraries were generated using the NEBNext Ultra II DNA Library Prep Kit (NEB, cat. no. E7645S) with 11 cycles of PCR amplification. Corresponding “Input” samples were processed identically. Two biological replicates were analyzed for each condition. ChIP-seq libraries analyzed in this study are listed in Table S4.

### MNase sensitivity analysis

Crosslinked mycelial powder (80 mg) was combined with 1 mL of sonication buffer containing 2 mM CaCl₂. Samples were equilibrated on ice for 10 min, mixed again, supplemented with 10 µL of MNase (NEB, cat. no. M0247S), and incubated in a ThermoMixer C (Eppendorf) for 30 min at 37°C. Reactions were stopped by adding 30 µL of 0.5 M EGTA (pH 8.0). Samples were cleared by centrifugation twice at 16,000 × g for 10 min at 4°C. Crosslink reversal and DNA purification were done as described in the ChIP protocol. Samples were resuspended in 30 µL of TE buffer. DNA concentration was measured on a Qubit using the dsDNA HS Assay kit (Invitrogen, cat. no. Q33230). Libraries were generated using the NEBNext Ultra II DNA Library Prep Kit (NEB, cat. no. E7645S) with 11 cycles of PCR amplification. MNase-seq libraries analyzed in this study are listed in Table S4.

### MNase-based ChIP

MNase treatment was performed as described above. Following quenching with EGTA and centrifugation, samples were processed for ChIP as described above, starting with addition of the appropriate antibody.

### Isolation of small RNAs for Illumina sequencing

Total RNA was extracted from mycelial powder using TRIzol reagent (Invitrogen, cat. no. 15596026) and dissolved in 5 M urea. RNA quality was assessed using an Agilent 4150 TapeStation system. Samples with RINe scores >9.5 were fractionated on 8% TBE-urea gels (60 µg of RNA per gel). Regions with small RNAs (sRNAs) in the 17-26 nt size range were excised with a razor blade, crushed, and incubated overnight in 0.3 M NaCl at 25°C with shaking at 850 rpm. Solution containing sRNAs was separated from gel debris using 0.22-µm spin columns (Sigma-Aldrich, cat. no. CLS8161). The filtered solution was divided into three 1.5-mL tubes; sRNAs were precipitated with 4 volumes of ethanol and pelleted by centrifugation at 20,000 × g for 40 min at 4°C. Pellets were resuspended in 4 µL of water each and combined to a total volume of 12 µL. sRNA-seq libraries were generated with 11 cycles of PCR amplification as previously described^9^. All sRNA-seq libraries analyzed in this study are listed in Table S4.

### Processing and analysis of Illumina sequencing data

#### A. Sequencing and post-processing

Sets of 48 libraries were sequenced on a NextSeq 500 or NextSeq 2000 platform (Illumina) in a single-end mode at the Institut Pasteur, Paris. Raw sequencing data were demultiplexed and converted to FASTQ files with bcl2fastq (version 2.20). Cutadapt^46^ (version 1.15) was used for adapter trimming.

#### B. Data analysis

Two principal reference genomes were used. Reference “T838” was used for strains carrying *tetO::ntcR* and *tetR-gfp*, whereas reference “T952” was used for strains carrying the unmodified *tetO* array and no *tetR-gfp*. Reference genomes are provided in Dataset 1. Reads were aligned using bowtie2^47^ as previously described^9^. Aligned data were analyzed using SAMtools^48^, BCFtools^48^, BEDTools^49^, deepTools^50^, and custom Python, Perl, and R scripts. ChIP-seq signal was computed as the number of reads per interval normalized to the total coverage equivalent of one million 75-nt reads. sRNA-seq signal was computed as the number of reads per interval normalized to the total number of aligned reads. H4v peaks were identified with MACS3^26^ (version 3.0.3) using the following parameters “-g 4.2e7-s 75-p 0.05 --broad-cutoff 0.01 --broad --nomodel --keep-dup all”. Peaks were classified into “AT-rich” or “GC-norm” based on the GC threshold of 41%, computed over the peak body and the 1000-bp flanking regions. Plots were generated using matplotlib^51^ and ggplot2^52^.

### Analysis of cytosine methylation by Nanopore sequencing

#### A. Library preparation and sequencing

Sequencing libraries were prepared using the Native Barcoding Kit 24 V14 (Oxford Nanopore Technologies) and sequenced on MinION flow cells according to the manufacturer’s instructions.

#### B. Data analysis

Basecalling was performed using Dorado version 0.9.1 (Oxford Nanopore Technologies) with the SUP model (version 4.2.0) and the 5mC modified-base model (version 2). Samples were demultiplexed with Dorado; reads from different sequencing runs were merged with SAMtools^48^ and aligned to the reference genome (T952) with Dorado. Summary alignment statistics were generated with NanoPlot^53^, SAMtools, and a custom R script. Methylated sites were identified using the pileup command of Modkit (Oxford Nanopore Technologies). A Hidden Markov model, implemented in the R package METHimpute^54^ (version 1.8.0), was used to classify each cytosine into either methylated or unmethylated. Modkit output was first converted into a METHimpute-compatible input format using a custom R script. Methylation profiles were visualized using ggplot2^52^.

### Fast Protein Liquid Chromatography (FPLC)

#### A. Preparation of cleared cell lysates

Mycelial powder (∼200 mg) was combined with 2 mL of sonication buffer (described above) and sonicated on ice using a Vibracell 75043 instrument (Bioblock Scientific) with the following program: total time = 2 min, pulse on = 2 s, pulse off = 8 s, 40% amplitude. Lysates were cleared by centrifugation twice at 20,000 × g for 5 min at 4°C.

#### B. Gel filtration chromatography

Chromatography was performed at ambient temperature using an ÄKTA Pure system (Cytiva) fitted with a Superdex 200 Increase 10/300 GL column (Cytiva, cat. no. 28990944). The column was equilibrated in SEC buffer (20 mM Tris-HCl, pH 8.0, 0.5 M NaCl). Cleared cell lysate (0.5 mL) was loaded onto the column, and separation was carried out with a flow rate of 0.5 mL/min. Elution was monitored by absorbance at 280 nm, and fractions of 0.8 mL were collected. The column was calibrated using standard protein markers (Sigma-Aldrich, cat. no. MWGF200).

#### C. Anion-exchange chromatography

Chromatography was performed at ambient temperature using an ÄKTA Pure system (Cytiva) fitted with a 1-mL HiTrap Q column (Cytiva, cat. no. 29051325). Two buffers were used to generate the elution gradient: buffer A (20 mM Tris-HCl, pH 8.0) and buffer B (20 mM Tris-HCl, pH 8.0, 1.0 M NaCl). Cleared lysate (2 mL) was loaded onto the column equilibrated with sonication buffer (described above). The column was then washed with 15 mL of 30% buffer B. Elution was performed with a custom gradient of buffer B (from 30% to 100%) at a flow rate of 0.5 mL/min. Elution was monitored by absorbance at 280 nm, and fractions of 0.7 mL were collected.

### Protein co-immunoprecipitation (co-IP)

Anti-FLAG M2 Magnetic Beads (Sigma-Aldrich, cat. no. M8823) were used. Beads (15 µL of suspension) were washed twice in 1 mL of TTBS (20 mM Tris-HCl, pH 7.5, 150 mM NaCl, 0.2% Tween-20) and combined with 1 mL of input material containing FLAG-tagged proteins. Samples were incubated overnight at 4°C on a rotating wheel. Subsequently, beads were washed three times with TTBS (each wash included a 10-min incubation at 4°C on a rotating wheel). Proteins were eluted with 50 µL of TES + NaCl buffer (50 mM Tris-HCl, pH 8.0, 10 mM EDTA, 1% SDS, 100 mM NaCl) and incubation at 50°C for 10 min at 1000 rpm in a ThermoMixer C (Eppendorf).

### Western blot analysis

Custom SDS-PAGE gels were made from 40% acrylamide/bis-acrylamide solution (37.5:1; Bio-Rad, cat. no. 1610148). Proteins were transferred onto Hybond P 0.2-µm PVDF membranes (Cytiva; cat. no. 10600021) using a Trans-Blot Turbo Transfer System (Bio-Rad). Membranes were blocked in 3% non-fat milk in TTBS for 30 min at ambient temperature and incubated with primary antibodies (1:5000 dilution) overnight at 4°C on a rocking platform. Membranes were washed three times with TTBS (each time for 10 min at 4°C) and incubated in TTBS with HRP-conjugated secondary antibody and 3% milk for 4 h at ambient temperature on a rocking platform. After three additional 10-min TTBS washes, protein signals were detected using Clarity Western ECL Substrate (Bio-Rad, cat. no. 1705060) and visualized using an Amersham Imager 680 system.

### Mass spectrometry analysis

#### A. Protein digestion and peptide cleanup

Proteins were denatured by adding 8 M urea in 100 mM Tris-HCl (pH 7.5). Proteins were reduced with 5 mM TCEP for 30 min at ambient temperature. Alkylation of reduced cysteine residues was performed using 20 mM iodoacetamide for 30 min at ambient temperature in the dark. Urea was diluted below 1.5 M with 50 mM ammonium bicarbonate. Sequencing-grade modified trypsin was added at an enzyme/protein ratio of 1:80 (w/w) in 100 mM ammonium bicarbonate, and samples were digested overnight at 37°C. Digestion was stopped by adding 1% formic acid. Samples were centrifuged at 16,000 × g for 10 min and the supernatant containing peptides was collected. Peptide clean-up was performed using an Agilent Bravo system with C18 StageTips according to the manufacturer’s protocol. Peptides were resuspended in 2% acetonitrile/0.1% formic acid prior to LC-MS injection.

#### B. Liquid chromatography-mass spectrometry

Peptides were analyzed by nano-liquid chromatography-tandem mass spectrometry (LC-MS/MS) using an EASY-nLC 1200 system (Thermo Fisher Scientific) coupled to an Orbitrap Q Exactive Plus mass spectrometer (Thermo Fisher Scientific). One µg of peptides was injected onto a PepMap RSLC 50 cm C18 column (2 µm particles, 100 Å pore size). Column equilibration and peptide loading were performed at 900 bar in buffer A (0.1% formic acid). Peptides were separated using a multi-step gradient of buffer B (80% acetonitrile, 0.1% formic acid): 3-9% for 5 min, 9-29% for 70 min, 29-56% for 30 min, 56-100% for 5 min, followed by 100% B for 7 min and re-equilibration at 3% B for 15 min, at a flow rate of 250 nL/min. Column temperature was maintained at 60°C. MS data were acquired using Xcalibur in data-dependent acquisition mode. MS1 scans were acquired at a resolution of 70,000 and MS2 scans (fixed first mass 100 m/z) at a resolution of 17,500. The AGC targets and maximum injection times were set to 3 × 10⁶ and 20 ms for MS1, and 1 × 10⁶ and 60 ms for MS2. The 10 most intense precursor ions were selected for fragmentation (Top 10) with a dynamic exclusion of 45 s. The isolation window was set to 1.6 m/z and normalized collision energy was set to 28 for HCD fragmentation. An underfill ratio of 1.0% (intensity threshold 1.7 × 10⁵) was used. Unassigned precursor ions and charge states 1, 7, 8, and >8 were excluded, and peptide match was disabled.

#### C. Data analysis

Data were analyzed with MaxQuant^55^ version 2.1.4 and the Andromeda search engine^56^. MS/MS spectra were searched against the *N. crassa* reference proteome provided by UniProt Knowledgebase (10,258 entries). The following modifications were considered: methionine oxidation, protein N-terminal acetylation, mono-, di-and trimethylation, GlyGly remnant (K), S/T/Y phosphorylation, and K acetylation; carbamidomethylation of cysteine was set as a fixed modification. Trypsin was specified as the protease, allowing up to two missed cleavages. The minimum peptide length was set to five amino acids, and the maximum peptide mass was 8,000 Da. The false discovery rate (FDR) was set to 1% at both peptide and protein levels. The main search tolerance was set to 4.5 ppm and MS/MS tolerance to 20 ppm. Second peptides were enabled. A minimum of one unique peptide was required for protein identification. MaxQuant output tables with protein, peptide, and PTM information are available as Dataset 4.

### Fluorescence microscopy analysis

Cultures were started by inoculating thawed macroconidia (∼10⁵ CFUs) into 5 mL of VM medium and grown for 6-7 h at 30°C with shaking at 160 rpm. 1 mL samples were collected and centrifuged at 10,000 × g, and mycelial pellets were mounted on glass slides. Images were acquired at ambient temperature using an inverted wide-field microscope (Nikon TE2000) equipped with a 100×/1.4 NA oil immersion objective, the adapted filter sets to observe green fluorescence (GFP-B Nikon: BP470/40, DC500, BP535/50), and a Prime BSI Express sCMOS camera (Teledyne Photometrics, 6.5 µm pixel size), controlled by MetaMorph software (Molecular Devices, RRID: SCR_002368). Samples were illuminated using a SPECTRA X Light Engine (Lumencor) at 50% power with an exposure time of 100 ms. For each experiment, all images were collected using the same acquisition settings and converted to 8-bit images using the same parameters. The images shown represent maximum intensity projections of 32-image z-stacks acquired with a 200 nm step. Raw images are available upon request.

### Luciferase-reporter assay to assess circadian rhythms

Macroconidia were collected from 7-day-old agar slants and inoculated into black 96-well plates containing the following medium: 1× Vogel’s salts, 0.1% glucose, 0.17% arginine, 1.5% agar, 50 ng/mL biotin, and 25 µM D-luciferin (GoldBio, cat. no. LUCK-10G). Plates were sealed with a Breathe-Easy sealing membrane (USA Scientific) and entrained in 12:12 light:dark cycles for 2 days, followed by incubation in constant darkness at 25°C for recording. Luciferase signals were recorded every hour (15 min exposure) for 5 days using a CCD camera (Pixis 1024B, Princeton Instruments) controlled by LightField software (Princeton Instruments, version 6.10.1). The mean intensity of each well was quantified and background-subtracted with an ImageJ^57^ macro^33^. Luminescence traces and period estimates were based on 3-4 technical replicates per each isolate. Circadian periods were estimated using a custom R script^58^. Statistical significance of period differences was assessed by one-way ANOVA followed by multiple-comparison testing using GraphPad Prism version 10.4.1 for macOS.

### Analysis of repeat-induced point mutation (RIP)

Crosses were set up in 90-mm Petri dishes (Gosselin, cat. no. SB90-101) using Synthetic Cross medium (1× SC salts, 2% sucrose, 0.2 µg/mL biotin, and 2% agar)^59^. Lids were replaced on day 13 post-fertilization. On day 24 post-fertilization, ejected ascospores were collected from the lids into water, heat-activated (30 min at 60°C), and plated on sorbose agar at appropriate dilutions. Cyclosporin-resistant colonies were picked with glass Pasteur pipettes and used to inoculate 2-mL cultures for genomic DNA extraction. The reporter region was amplified by PCR using Phusion Hot Start DNA Polymerase (NEB, cat. no. M0535S) and analyzed by Sanger sequencing at Eurofins Genomics (Cologne, Germany). Chromatograms were assembled into contigs using Phred/Phrap^60^; assemblies were inspected in Consed^61^. Contigs were aligned to the reference sequence using ClustalX^62^. The occurrence of mutations was determined using a custom Perl script. Analyzed crosses are listed in Table S5. Sequence alignments are provided in Dataset 5.

### Prediction of H4v-containing nucleosome structure

A structural model of a nucleosome containing two copies of H4v was created using the AlphaFold3^63^ server (https://alphafoldserver.com). The input consisted of eight polypeptide chains corresponding to two copies of each *Neurospora* histone (H2A, H2B, H3, and H4v) and two complementary DNA strands corresponding to the Widom 601 positioning sequence^64^. Structure prediction was performed without applying constraints on the histone octamer. The resulting model was visualized in PyMOL (Schrödinger). Five alternative predicted nucleosome structures are provided in Dataset 3.

## SUPPLEMENTARY FIGURE LEGENDS

**Figure S1.**
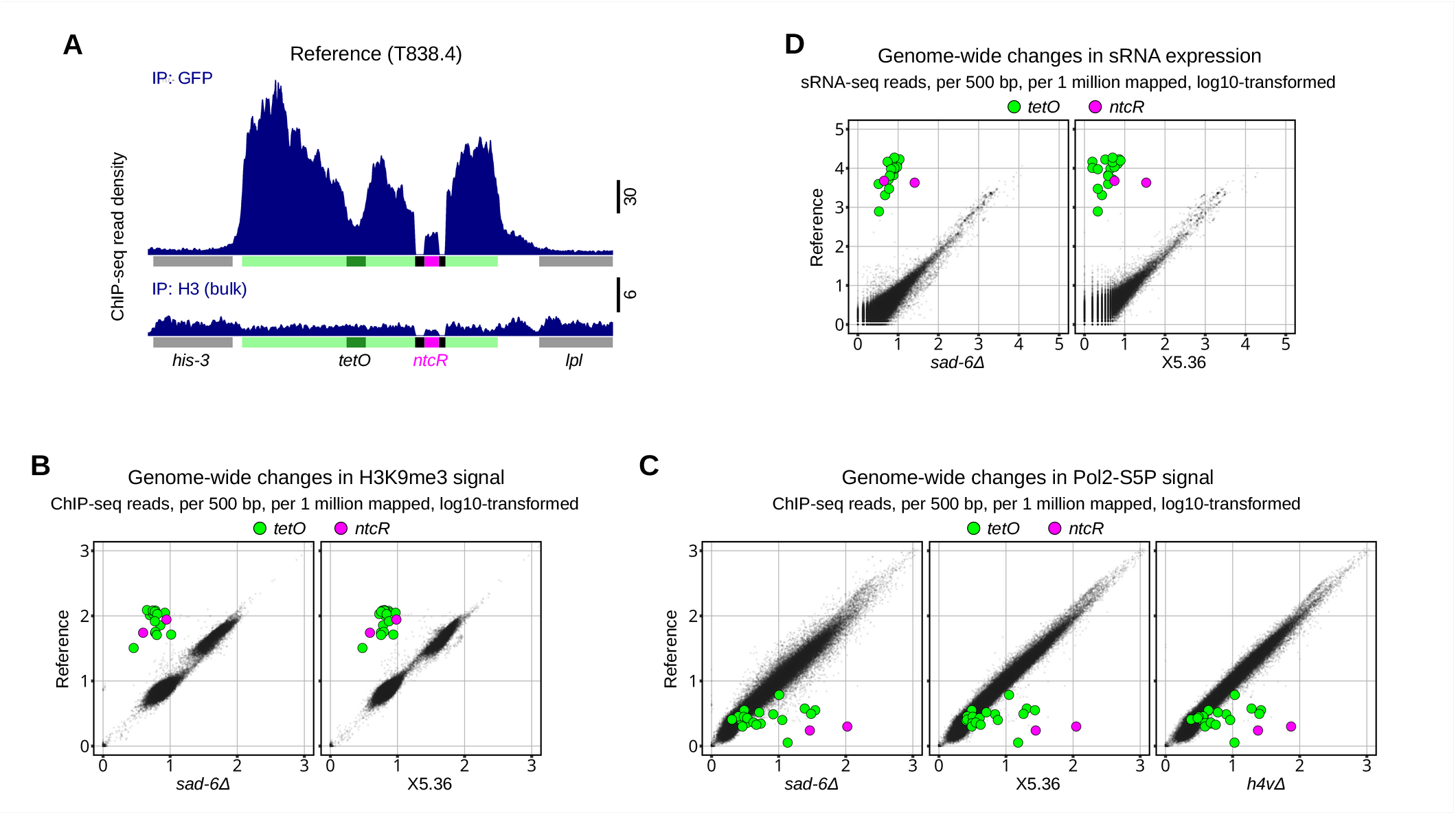
Genome-wide effects of *sad-6Δ* and *h4vΔ* compared to isolate X5.36. **(A)** GFP and bulk H3 ChIP-seq profiles at the *tetO::ntcR* locus in strain T838.4 (Reference). **(B)** Genome-wide comparison of H3K9me3 ChIP-seq profiles between indicated genetic backgrounds, analyzed as in Fig. 1E. Analyzed strains are listed in Fig. 1G. **(C)** Genome-wide comparison of Pol2-S5P ChIP-seq profiles between indicated genetic backgrounds, analyzed as in Fig. 1E. Analyzed strains are listed in Fig. 1G. **(D)** Genome-wide comparison of small RNA production between indicated genetic backgrounds, analyzed as in Fig. 1F. Analyzed strains are listed in Fig. 1G.

**Figure S2.**
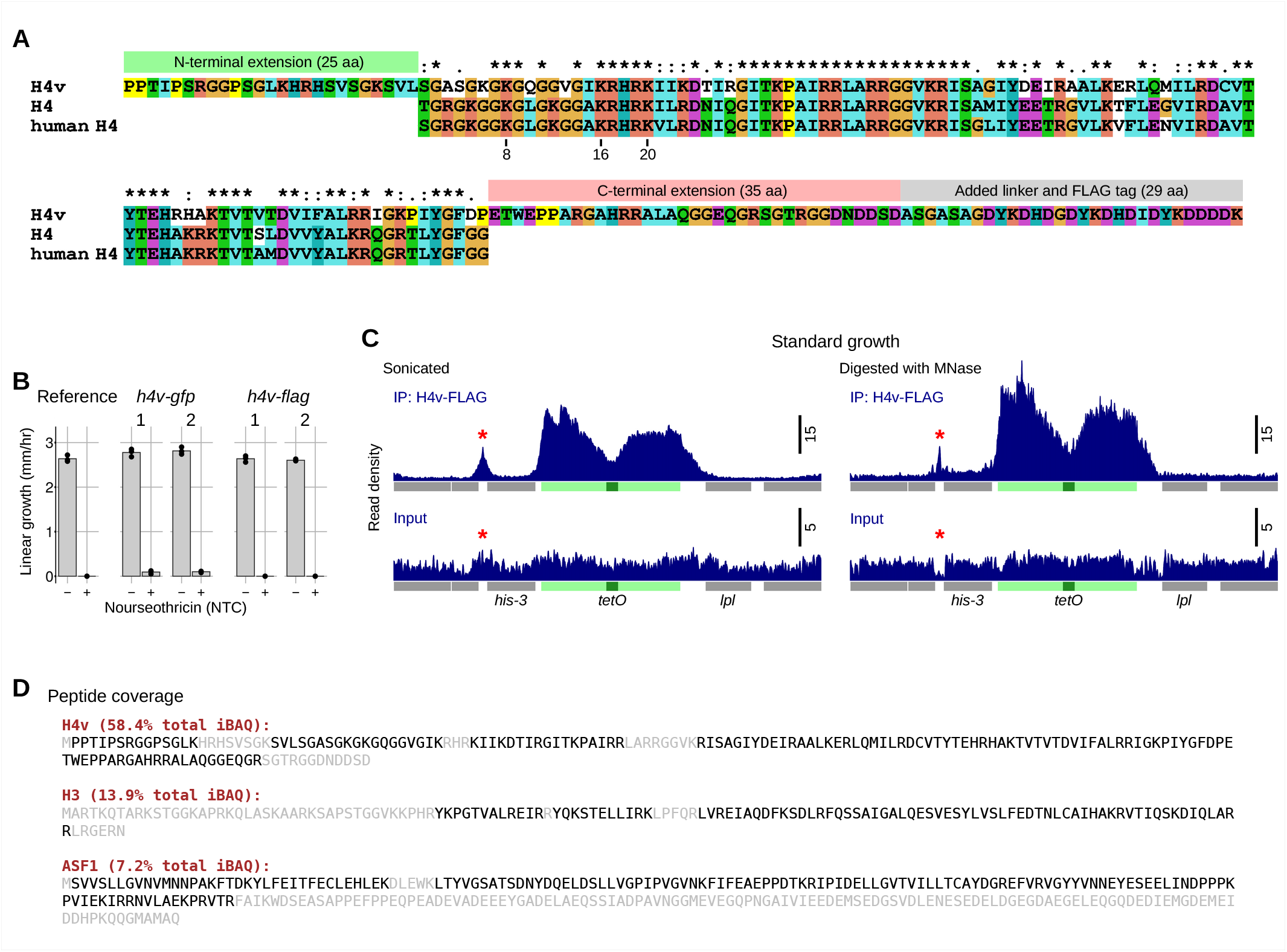
H4v sequence features, effects of C-terminal tags, MNase-based ChIP-seq validation, and identification of H4v-associated proteins. **(A)** Amino acid sequence alignment of *Neurospora* H4v, canonical H4, and human H4. The H4v-specific N-terminal extension (25 aa), unique C-terminal part (35 aa), and appended FLAG tag are indicated. Conserved residues are marked; canonical H4 numbering is shown above the alignment. **(B)** Linear growth of *N. crassa* strains in the presence/absence of NTC. The following strains were analyzed: T838.4 (Reference, data also reported in Fig. 1D); T930.5h and T930.8h (*h4v-flag*); T962.4h and T962.6h (*h4v-gfp*). **(C)** H4v-FLAG ChIP-seq profiles at the *tetO* locus obtained from sonicated (left) and MNase-digested (right) chromatin. Asterisk marks an MNase-sensitive region retaining H4v. **(D)** Peptide coverage for H4v, H3, and ASF1 identified by mass spectrometry of the H4v-FLAG immunoprecipitate from Superdex fraction 8 (Fig. 3A). Detected peptides are highlighted. For each protein, the summed intensity-based absolute quantification (iBAQ) value is expressed as a percentage of the total iBAQ signal.

**Figure S3.**
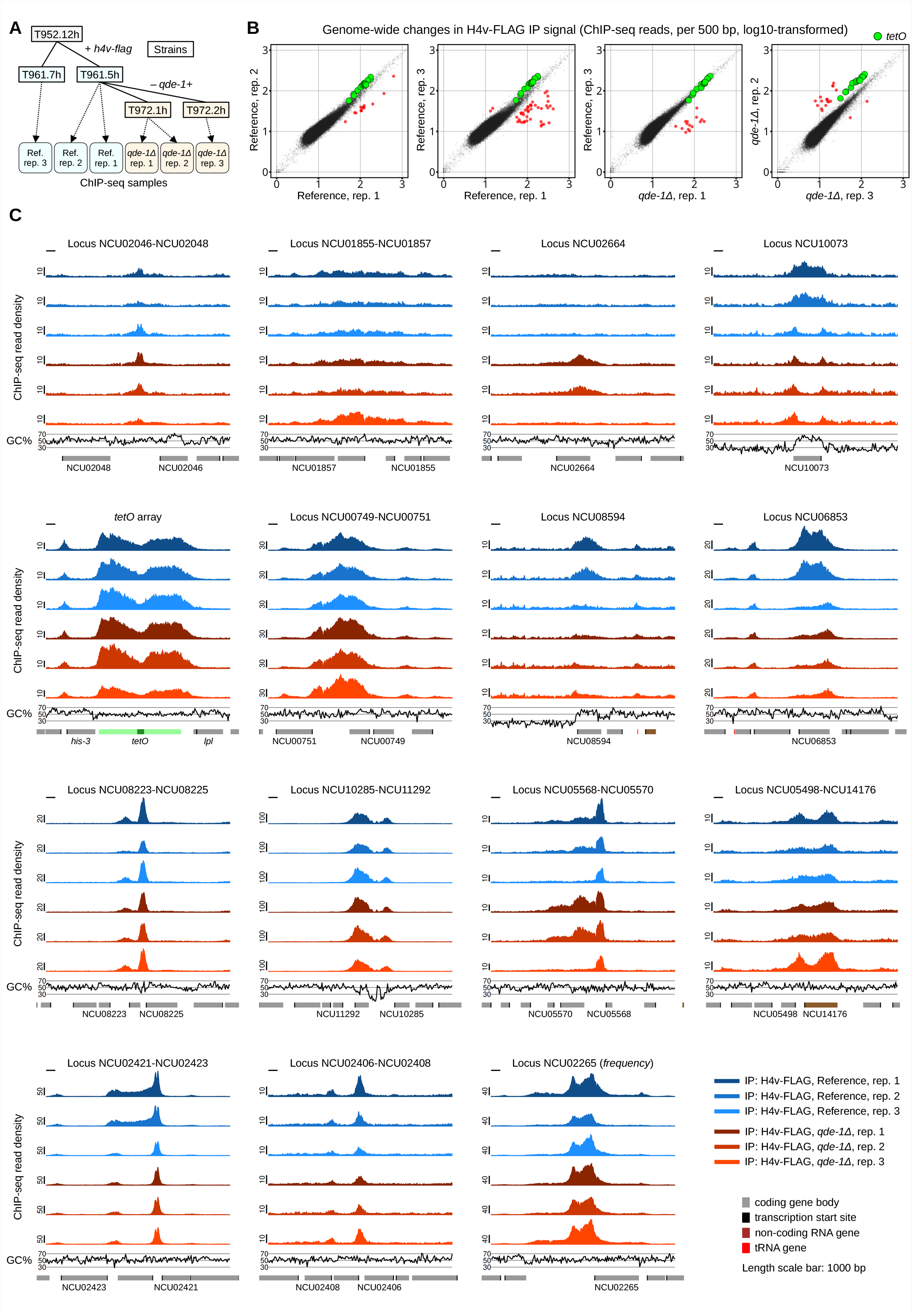
Several loci exhibit highly variable H4v-FLAG ChIP-seq signal. **(A)** Schematic showing the relationships among the analyzed ChIP-seq samples, including biological and technical replicates. **(B)** Representative genome-wide comparisons of H4v-FLAG ChIP-seq profiles between several samples in Fig. S3A. Analyzed as in Fig. 1E. **(C)** H4v-FLAG ChIP-seq profiles for loci with highly variable H4v enrichment (corresponding 500-bp bins are shown as red circles in Fig. S3B). GC content and gene annotations are shown below. Scale bar, 1000 bp.

**Figure S4.**
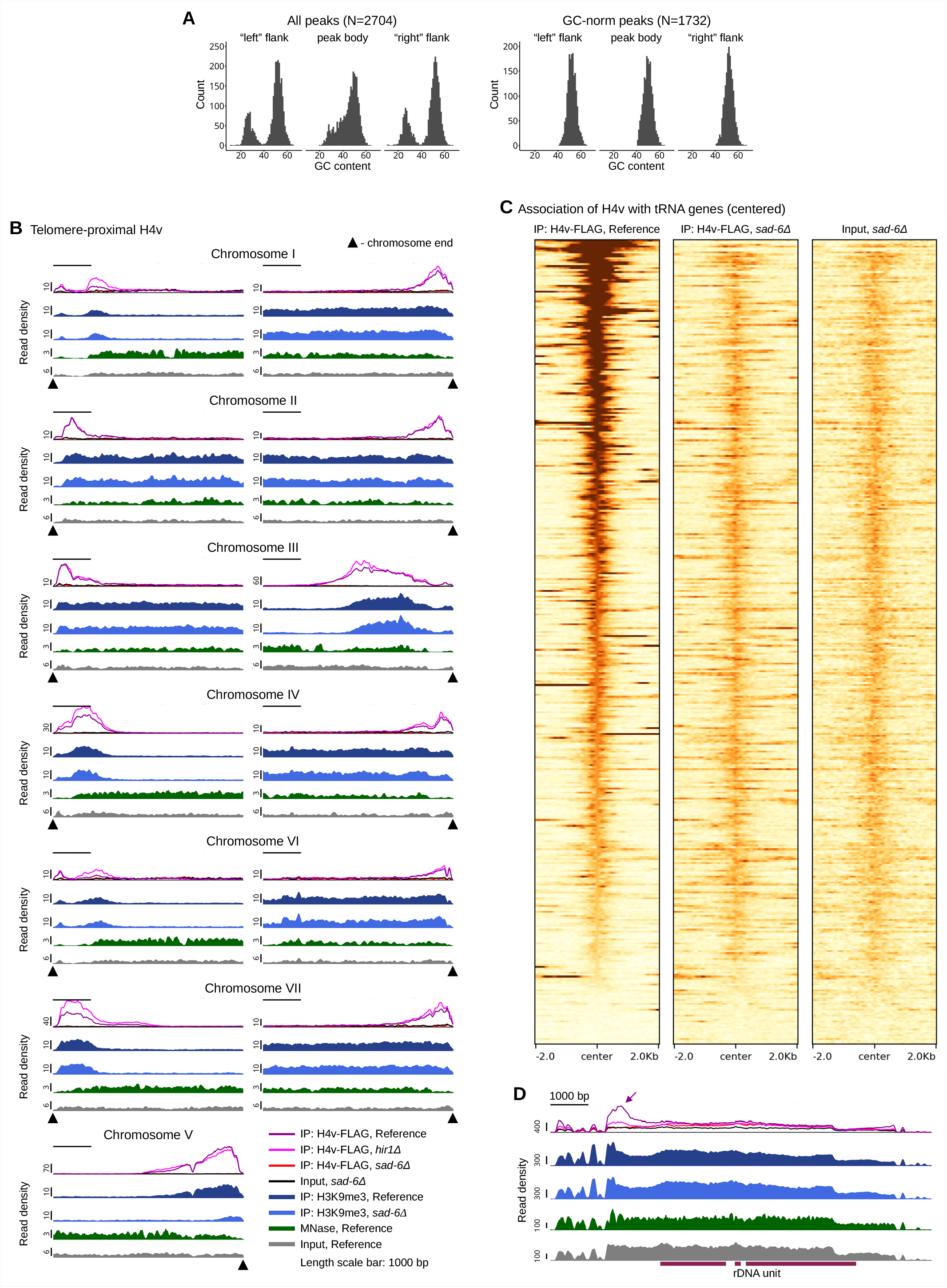
Classification of H4v peaks based on GC context, H4v enrichment at telomeres, rDNA, and tRNA genes. **(A)** GC content distributions in peak flanks and bodies for all peaks (N = 2,704; left) and GC-norm peaks only (N = 1,732; right). **(B)** ChIP-seq profiles at telomere-proximal regions across all seven chromosomes. Tracks show H4v-FLAG (Reference, *hir1Δ*, *sad-6Δ*), H3K9me3 (Reference, *sad-6Δ*), MNase (Reference), and Input levels (Reference and *sad-6Δ*). Analyzed strains are listed in Fig. 2E. Triangles mark chromosome ends. Scale bar, 1000 bp. **(C)** Heatmaps of H4v-FLAG ChIP-seq signal (Reference and *sad-6Δ*) centered on the annotated tRNA genes (±2 kb). Input is provided for comparison. Each row represents one tRNA gene. **(D)** ChIP-seq profiles across the rDNA repeat unit in the reference, *hir1Δ*, and *sad-6Δ* backgrounds together with the reference Input levels. The coloring scheme is provided in Fig. S4B. Scale bar, 1,000 bp.

**Figure S5.**
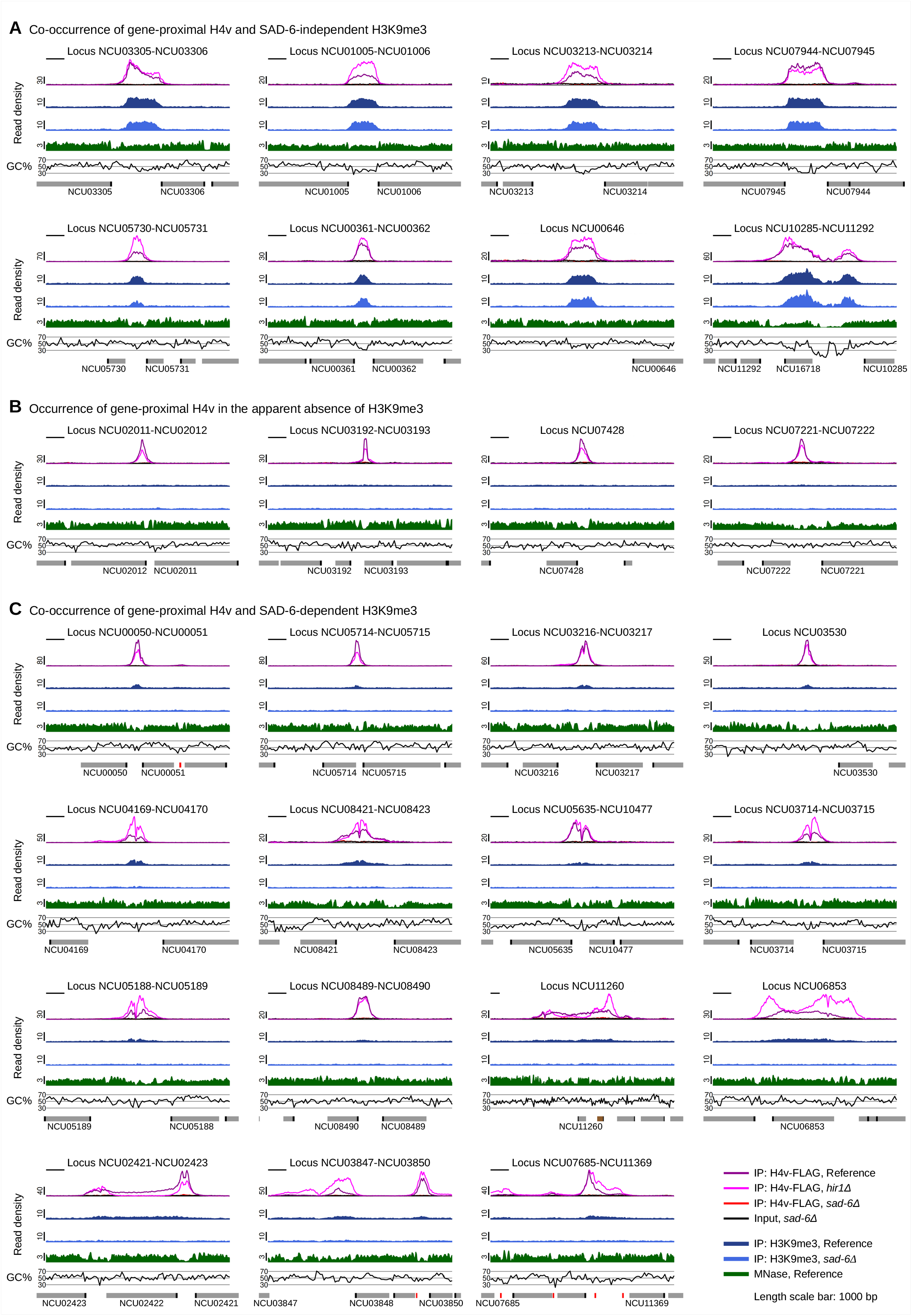
Three classes of gene-proximal H4v peaks defined by H3K9me3 enrichment. For all panels, tracks show H4v-FLAG (Reference, *hir1Δ*, *sad-6Δ*), H3K9me3 (Reference, *sad-6Δ*), MNase (Reference), and Input (*sad-6Δ*). GC content and gene annotations are plotted below. Analyzed strains are listed in Fig. 2E. Scale bar, 1000 bp. **(A)** H4v peaks with strong, SAD-6-independent H3K9me3, associated with local reductions in GC content. **(B)** H4v peaks without detectable H3K9me3. **(C)** H4v peaks with weak, SAD-6-dependent H3K9me3 (including known *disiRNA* loci).

**Figure S6.**
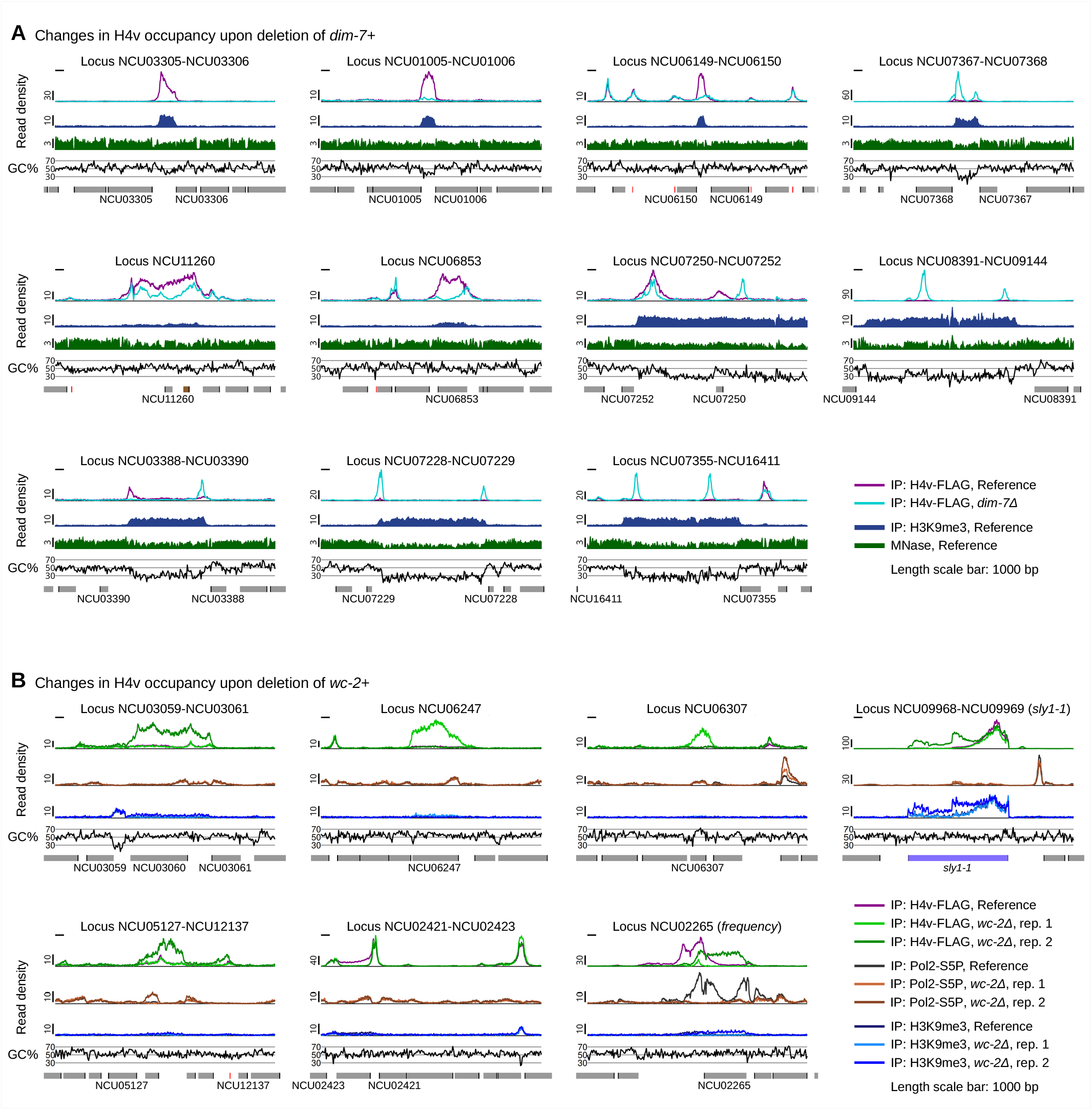
Changes in H4v occupancy upon deletion of *dim-7+* or *wc-2+*. **(A)** H4v-FLAG and H3K9me3 ChIP-seq profiles at selected loci in the reference and *dim-7Δ* backgrounds. Tracks show H4v-FLAG (Reference, *dim-7Δ*), H3K9me3 (Reference), and MNase (Reference). GC content is plotted below. Analyzed strains are listed in Fig. 5A. Scale bar, 1000 bp. **(B)** H4v-FLAG, Pol2-S5P, and H3K9me3 ChIP-seq profiles at selected loci in the reference and *wc-2Δ* backgrounds. Analyzed strains are listed in Fig. 5A. Scale bar, 1000 bp.

**Figure S7.**
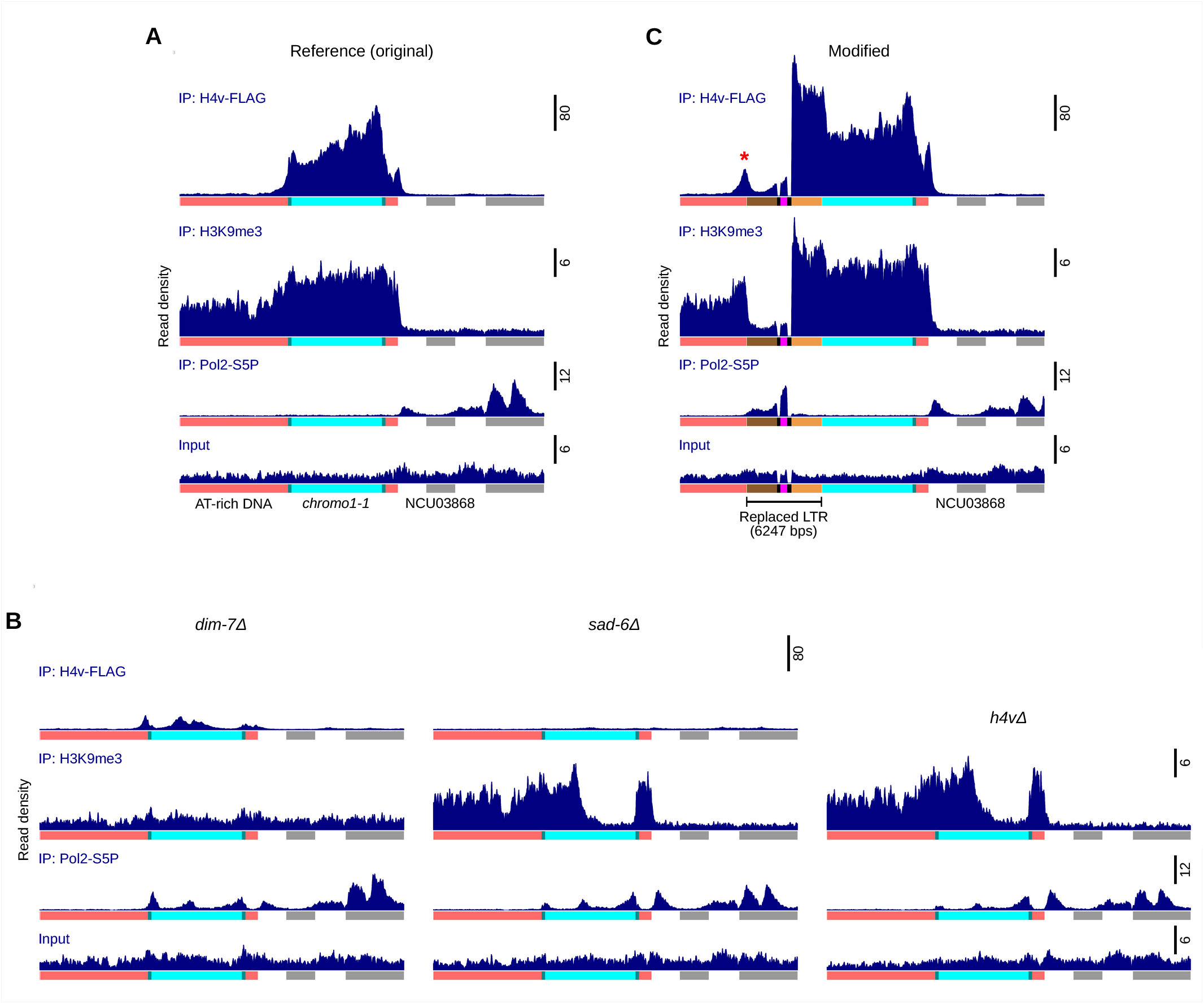
Chromatin profiles at the wild-type and modified *chromo1-1* element. **(A)** H4v-FLAG, H3K9me3, and Pol2-S5P ChIP-seq profiles along with the corresponding Input levels for the *chromo1-1* locus in the reference background. The analyzed strains are listed in Fig. 2E. **(B)** Same profiles as in Fig. S7A for *dim-7Δ*, *sad-6Δ*, and *h4vΔ* backgrounds. The analyzed *dim-7Δ* strains are listed in Fig. 5A; the *sad-6Δ* strains are listed in Fig. 2E; the *h4vΔ* strains were the following: T979.1h and T979.2h. **(C)** Same profiles as in Fig. S7A for the modified *chromo1-1* in which the “left” LTR was replaced with a 6,247-bp construct containing *ntcR* flanked by *E. coli* DNA. The replaced region is indicated. An asterisk marks a H4v peak emerged at the AT-rich/*E. coli* DNA boundary. The following strains were analyzed: T1035.1h and T1035.2h.

**Figure S8.**
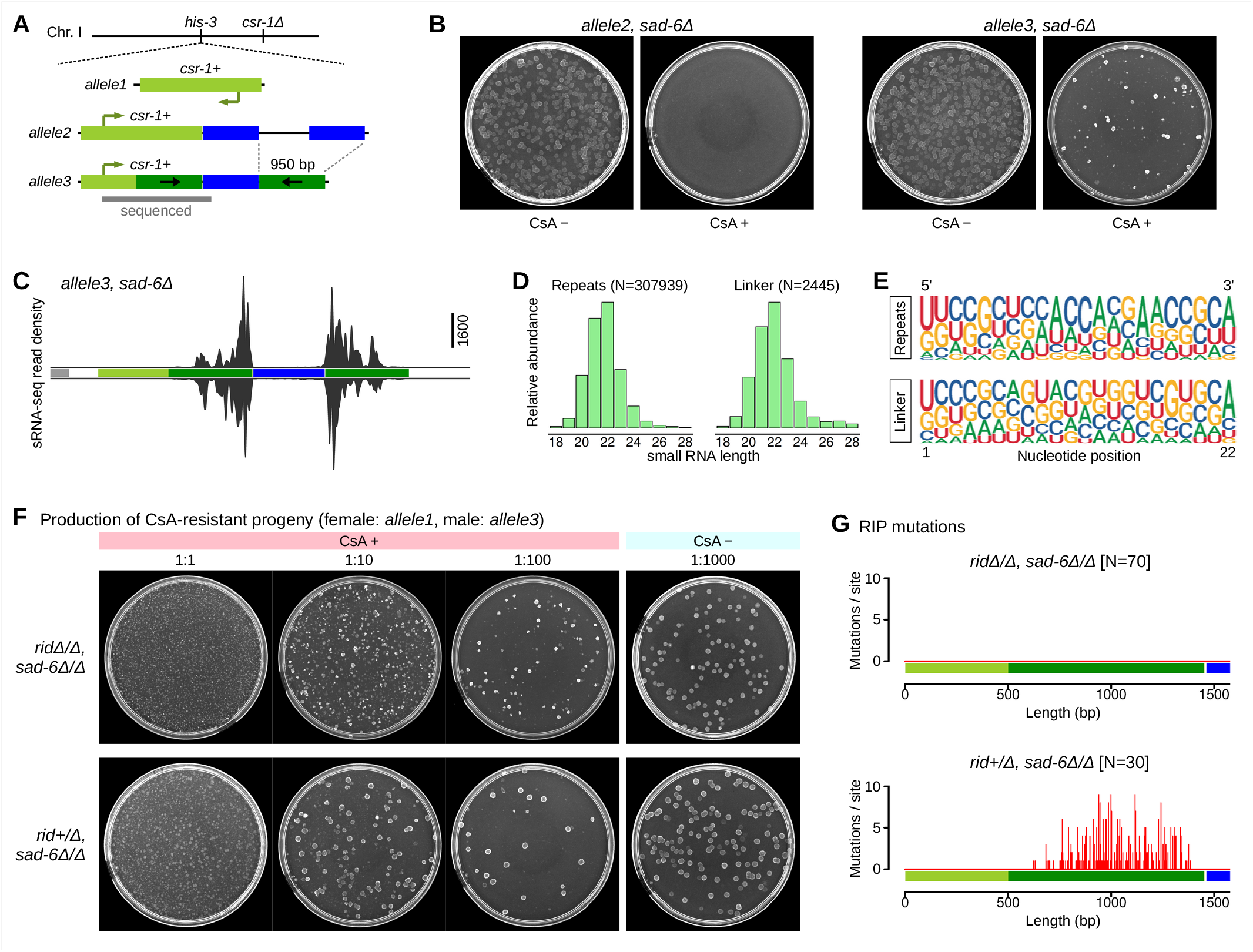
SAD-6 is dispensable for RID-dependent RIP. **(A)** Additional reporter (*allele3*) was generated to provide a more efficient substrate for RID-dependent RIP at the ectopic *csr-1+* locus. Gray bar denotes a region analyzed by sequencing in Fig. S8G. **(B)** Growth of *sad-6Δ* strains containing *allele2* (strain T1000.2h) or *allele3* (strain T1033.3h) in the absence (CsA−) and presence (CsA+) of cyclosporin A. **(C)** sRNA-seq profile of *allele3* in the *sad-6Δ* strain T1033.3h. **(D)** Size distribution of small RNAs mapping to the repeated segments (N = 307,939) and linker (N = 2,445) of the *allele3* construct. **(E)** Nucleotide composition logos for small RNAs from the repeated segments (top) and linker (bottom). **(F)** Production of CsA-resistant progeny from crosses carrying *allele3* in the indicated genetic backgrounds. Serial dilutions were analyzed as in Fig. 6D. Crosses between the following strains were analyzed (listed as female x male parents): T996.5h x T1033.3h and T996.5h x T1033.8h (*ridΔ/Δ, sad-6Δ/Δ*); T1037.3h x T1033.3h and T1037.4h x T1033.8h (*rid+/Δ, sad-6Δ/Δ*). Analyzed crosses are listed in Table S5. **(G)** Distribution of RIP mutations across the reporter locus in the indicated genetic backgrounds. Mutation frequency is plotted as a function of position along the repeat construct. Analyzed multiple sequence alignments are provided in Dataset 5, referenced by cross names listed in Table S5.

## SUPPLEMENTARY DATASETS

**Dataset 1. Reference genomes (in FASTA format).**

**Dataset 2. Transformation plasmids (in EMBL format).**

**Dataset 3. AlphaFold3 models of the H4v-containing nucleosome (in mmCIF format).**

**Dataset 4. MaxQuant-generated proteomics results (in Excel format).**

**Dataset 5. Multiple sequence alignments analyzed for RIP (in CLUSTAL format).**

All datasets are available at Figshare (DOI: 10.6084/m9.figshare.32952437).

---

**Table S1.** *N. crassa* genes analyzed in this study.

| Standard name | Gene ID | Genome accession | Start position | End position | Protein accession | Full name | Synonym / ortholog |
| --- | --- | --- | --- | --- | --- | --- | --- |
| *asf1* | NCU09436 | NC_026507.1 | 178902 | 181174 | XP_958588.1 | *anti-silencing function 1* |  |
| *csr-1* | NCU00726 | NC_026501.1 | 7403946 | 7406381 | XP_011392822.1 | *cyclosporin-resistant-1* | cyp; CyP20 |
| *dim-2* | NCU02247 | NC_026507.1 | 3189797 | 3195483 | XP_959891.1 | *defective in methylation-2* |  |
| *dim-5* | NCU04402 | NC_026504.1 | 3480589 | 3482273 | XP_957479.2 | *defective in methylation-5* | Su(var)3-9 |
| *dim-7* | NCU04152 | NC_026505.1 | 5011283 | 5014974 | XP_961308.2 | *defective in methylation-7* |  |
| *eri1* | NCU06684 | NC_026505.1 | 4263047 | 4267333 | XP_011394691.1 | *exoribonuclease 1* |  |
| *frq* | NCU02265 | NC_026507.1 | 3131640 | 3136464 | XP_011395125.1 | *frequency* |  |
| *hh3* | NCU01635 | NC_026502.1 | 2874241 | 2875647 | XP_956003.1 | *histone H3* |  |
| *hh4-1* | NCU01634 | NC_026502.1 | 2877350 | 2878740 | XP_956002.2 | *histone H4* |  |
| *hh4-2* | NCU00212 | NC_026503.1 | 3024058 | 3025897 | XP_956597.1 | *histone H4* |  |
| *hh4v* | NCU04338 | NC_026504.1 | 3729161 | 3731640 | XP_956319.3 | *histone H4 variant* | H4v; H4E |
| *hir1* | NCU04035 | NC_026506.1 | 2519404 | 2524121 | XP_957650.2 | *histone regulation 1* |  |
| *his-3* | NCU03139 | NC_026501.1 | 4694770 | 4697885 | XP_964188.1 | *histidine-3* |  |
| *lpl* | NCU03141 | NC_026501.1 | 4700673 | 4703568 | XP_964190.2 | *lysophospholipase* |  |
| *mus-52* | NCU00077 | NC_026503.1 | 2531930 | 2535354 | XP_956387.3 | *mutagen sensitive-52* | KU80 |
| *qde-1* | NCU07534 | NC_026503.1 | 4461937 | 4467476 | XP_959047.1 | *quelling-defective-1* |  |
| *rid* | NCU02034 | NC_026501.1 | 1611869 | 1615028 | XP_011392925.1 | *RIP defective* |  |
| *sad-6* | NCU06190 | NC_026503.1 | 1838943 | 1845994 | XP_963002.3 | *suppressor of ascus dominance-6* | ATRX; XNP |
| *set-7* | NCU07496 | NC_026501.1 | 408524 | 415263 | XP_965043.2 | *set-domain histone methyltransferase-7* | E(z) |
| *wc-2* | NCU00902 | NC_026501.1 | 6815517 | 6819124 | XP_963819.3 | *white collar-2* |  |

**Table S2.** *N. crassa* strains used in this study.

| Strain ID | Genotype | Source |
| --- | --- | --- |
| C02.1 | *A, ridΔ, mus-52Δ, his-3* | Ref. 18 |
| C03.1 | *a, ridΔ, mus-52Δ, his-3* | Ref. 18 |
| C139.10 | *a, set-7Δ, ridΔ, mus-52Δ, his-3* | Ref. 18 |
| T658.1 | *a, ridΔ, dim-2Δ, mus-52Δ, his-3+::lacO-tetO, csr-1Δ::tetR-gfp* | Ref. 9 |
| T707.51h | *a, ridΔ, dim-2Δ, mus-52Δ, his-3+::lacO-tetO, csr-1Δ::tetR-gfp, sad-6Δ* | Ref. 9 |
| T798.1 | *A, ridΔ, mus-52Δ, csr-1Δ, his-3* | C02.1 transformed with pEAG66 |
| T799.1 | *a, ridΔ, mus-52Δ, csr-1Δ, his-3* | C03.1 transformed with pEAG66 |
| T804.8h | *a, ridΔ, mus-52Δ, csr-1Δ, his-3+::csr-1+* | T799.1 transformed with pEAG82 |
| T805.1h | *A, ridΔ, mus-52Δ, csr-1Δ, his-3+::FOC100T* | T798.1 transformed with pFOC100T |
| T827.7h | *a, ridΔ, dim-2Δ, mus-52Δ, his-3+::lacO-tetO, csr-1Δ::tetR-gfp, flag-sad-6* | Ref. 9 |
| T830.14h | *a, set-7Δ, ridΔ, mus-52Δ, his-3+::tetO::ntcR* | C139.10 transformed with pFOC104C1 |
| T838.4 | *a, set-7Δ, ridΔ, mus-52Δ, his-3+::tetO::ntcR, csr-1Δ::tetR-gfp* | T830.14h transformed with pTSN7B |
| X5.36.M1 | *a, set-7Δ, ridΔ, mus-52Δ, his-3+::tetO::ntcR, csr-1Δ::tetR-gfp* | Mutant isolate of T838.4 |
| X5.36.M2 | *a, set-7Δ, ridΔ, mus-52Δ, his-3+::tetO::ntcR, csr-1Δ::tetR-gfp* | Mutant isolate of T838.4 |
| T902.1h | *a, set-7Δ, ridΔ, mus-52Δ, his-3+::tetO::ntcR, csr-1Δ::tetR-gfp, sad-6Δ* | T838.4 transformed with pJBN1 |
| T902.4h | *a, set-7Δ, ridΔ, mus-52Δ, his-3+::tetO::ntcR, csr-1Δ::tetR-gfp, sad-6Δ* | T838.4 transformed with pJBN1 |
| T914.1h | *a, set-7Δ, ridΔ, mus-52Δ, his-3+::tetO::ntcR, csr-1Δ::tetR-gfp, dim-5Δ* | T838.4 transformed with pJBN2B |
| T914.3h | *a, set-7Δ, ridΔ, mus-52Δ, his-3+::tetO::ntcR, csr-1Δ::tetR-gfp, dim-5Δ* | T838.4 transformed with pJBN2B |
| T922.1h | *a, set-7Δ, ridΔ, mus-52Δ, his-3+::tetO::ntcR, csr-1Δ::tetR-gfp, h4vΔ* | T838.4 transformed with pJBN10A |
| T922.4h | *a, set-7Δ, ridΔ, mus-52Δ, his-3+::tetO::ntcR, csr-1Δ::tetR-gfp, h4vΔ* | T838.4 transformed with pJBN10A |
| T922.6h | *a, set-7Δ, ridΔ, mus-52Δ, his-3+::tetO::ntcR, csr-1Δ::tetR-gfp, h4vΔ* | T838.4 transformed with pJBN10A |
| T923.1h | *a, set-7Δ, ridΔ, mus-52Δ, his-3+::tetO::ntcR, csr-1Δ::tetR-gfp, dim-2Δ* | T838.4 transformed with pJBN11B |
| T923.2h | *a, set-7Δ, ridΔ, mus-52Δ, his-3+::tetO::ntcR, csr-1Δ::tetR-gfp, dim-2Δ* | T838.4 transformed with pJBN11B |
| T930.5h | *a, set-7Δ, ridΔ, mus-52Δ, his-3+::tetO::ntcR, csr-1Δ::tetR-gfp, h4v-flag* | T838.4 transformed with pJBN10E |
| T930.8h | *a, set-7Δ, ridΔ, mus-52Δ, his-3+::tetO::ntcR, csr-1Δ::tetR-gfp, h4v-flag* | T838.4 transformed with pJBN10E |
| T948.1h | *a, ridΔ, dim-2Δ, mus-52Δ, his-3+::lacO-tetO, csr-1Δ::tetR-gfp, h4v-flag* | T658.1 transformed with pJBN10E |
| T948.7h | *a, ridΔ, dim-2Δ, mus-52Δ, his-3+::lacO-tetO, csr-1Δ::tetR-gfp, h4v-flag* | T658.1 transformed with pJBN10E |
| T949.1h | *a, ridΔ, dim-2Δ, mus-52Δ, his-3+::lacO-tetO, csr-1Δ::tetR-gfp, sad-6Δ, h4v-flag* | T707.51h transformed with pJBN10E |
| T949.2h | *a, ridΔ, dim-2Δ, mus-52Δ, his-3+::lacO-tetO, csr-1Δ::tetR-gfp, sad-6Δ, h4v-flag* | T707.51h transformed with pJBN10E |
| T950.4h | *a, ridΔ, dim-2Δ, mus-52Δ, his-3+::lacO-tetO, csr-1Δ::tetR-gfp, flag-sad-6, h4vΔ* | T827.7h transformed with pJBN10A |
| T950.6h | *a, ridΔ, dim-2Δ, mus-52Δ, his-3+::lacO-tetO, csr-1Δ::tetR-gfp, flag-sad-6, h4vΔ* | T827.7h transformed with pJBN10A |
| T952.12h | *a, set-7Δ, ridΔ, mus-52Δ, his-3+::tetO* | C139.10 transformed with pTSN6 |
| T959.1h | *a, set-7Δ, ridΔ, mus-52Δ, his-3+::tetO, h4v-gfp* | T952.12h transformed with pJBN10G |
| T961.5h | *a, set-7Δ, ridΔ, mus-52Δ, his-3+::tetO, h4v-flag* | T952.12h transformed with pJBN10E |
| T961.7h | *a, set-7Δ, ridΔ, mus-52Δ, his-3+::tetO, h4v-flag* | T952.12h transformed with pJBN10E |
| T962.4h | *a, set-7Δ, ridΔ, mus-52Δ, his-3+::tetO::ntcR, csr-1Δ::tetR-gfp, h4v-gfp* | T838.4 transformed with pJBN10G |
| T962.6h | *a, set-7Δ, ridΔ, mus-52Δ, his-3+::tetO::ntcR, csr-1Δ::tetR-gfp, h4v-gfp* | T838.4 transformed with pJBN10G |
| T964.1h | *a, set-7Δ, ridΔ, mus-52Δ, his-3+::tetO, h4v-gfp, sad-6Δ* | T959.1h transformed with pFOC109B |
| T965.4h | *a, set-7Δ, ridΔ, mus-52Δ, his-3+::tetO, h4v-flag, sad-6Δ* | T961.5h transformed with pFOC109B |
| T965.9h | *a, set-7Δ, ridΔ, mus-52Δ, his-3+::tetO, h4v-flag, sad-6Δ* | T961.5h transformed with pFOC109B |
| T965.12h | *a, set-7Δ, ridΔ, mus-52Δ, his-3+::tetO, h4v-flag, sad-6Δ* | T961.5h transformed with pFOC109B |
| T968.1h | *a, set-7Δ, ridΔ, mus-52Δ, his-3+::tetO, h4v-flag, hir1Δ* | T961.5h transformed with pFOC128B |
| T968.3h | *a, set-7Δ, ridΔ, mus-52Δ, his-3+::tetO, h4v-flag, hir1Δ* | T961.5h transformed with pFOC128B |
| T971.2h | *a, set-7Δ, ridΔ, mus-52Δ, his-3+::tetO, h4v-flag, dim-7Δ* | T961.5h transformed with pFOC116B |
| T971.3h | *a, set-7Δ, ridΔ, mus-52Δ, his-3+::tetO, h4v-flag, dim-7Δ* | T961.5h transformed with pFOC116B |
| T972.1h | *a, set-7Δ, ridΔ, mus-52Δ, his-3+::tetO, h4v-flag, qde-1Δ* | T961.5h transformed with pFOC120B |
| T972.2h | *a, set-7Δ, ridΔ, mus-52Δ, his-3+::tetO, h4v-flag, qde-1Δ* | T961.5h transformed with pFOC120B |
| T974.6h | *a, set-7Δ, ridΔ, mus-52Δ, his-3+::tetO, h4v-flag, eri1Δ* | T961.5h transformed with pSCR8B |
| T974.7h | *a, set-7Δ, ridΔ, mus-52Δ, his-3+::tetO, h4v-flag, eri1Δ* | T961.5h transformed with pSCR8B |
| T976.5h | *a, set-7Δ, ridΔ, mus-52Δ, his-3+::tetO, h4v-flag, asf1-ha* | T961.5h transformed with pEAG283F |
| T979.1h | *a, set-7Δ, ridΔ, mus-52Δ, his-3+::tetO, h4vΔ* | T952.12h transformed with pJBN10A |
| T979.2h | *a, set-7Δ, ridΔ, mus-52Δ, his-3+::tetO, h4vΔ* | T952.12h transformed with pJBN10A |
| T980.6h | *a, set-7Δ, ridΔ, mus-52Δ, his-3+::tetO, h4v-flag, wc-2Δ* | T961.5h transformed with pFOC146B |
| T980.12h | *a, set-7Δ, ridΔ, mus-52Δ, his-3+::tetO, h4v-flag, wc-2Δ* | T961.5h transformed with pFOC146B |
| T996.5h | *a, ridΔ, mus-52Δ, csr-1Δ, his-3+::csr-1+, sad-6Δ* | T804.8h transformed with pJBN1 |
| T996.8h | *a, ridΔ, mus-52Δ, csr-1Δ, his-3+::csr-1+, sad-6Δ* | T804.8h transformed with pJBN1 |
| T997.7h | *a, ridΔ, mus-52Δ, csr-1Δ, his-3+::csr-1+, h4vΔ* | T804.8h transformed with pJBN10A |
| T997.8h | *a, ridΔ, mus-52Δ, csr-1Δ, his-3+::csr-1+, h4vΔ* | T804.8h transformed with pJBN10A |
| T998.4h | *a, ridΔ, mus-52Δ, csr-1Δ, his-3+::csr-1+, dim-2Δ* | T804.8h transformed with pJBN11B |
| T998.6h | *a, ridΔ, mus-52Δ, csr-1Δ, his-3+::csr-1+, dim-2Δ* | T804.8h transformed with pJBN11B |
| T1000.2h | *A, ridΔ, mus-52Δ, csr-1Δ, his-3+::FOC100T, sad-6Δ* | T805.1h transformed with pJBN1 |
| T1000.12h | *A, ridΔ, mus-52Δ, csr-1Δ, his-3+::FOC100T, sad-6Δ* | T805.1h transformed with pJBN1 |
| T1001.3h | *A, ridΔ, mus-52Δ, csr-1Δ, his-3+::FOC100T, h4vΔ* | T805.1h transformed with pJBN10A |
| T1001.4h | *A, ridΔ, mus-52Δ, csr-1Δ, his-3+::FOC100T, h4vΔ* | T805.1h transformed with pJBN10A |
| T1002.2h | *A, ridΔ, mus-52Δ, csr-1Δ, his-3+::FOC100T, dim-2Δ* | T805.1h transformed with pJBN11B |
| T1002.5h | *A, ridΔ, mus-52Δ, csr-1Δ, his-3+::FOC100T, dim-2Δ* | T805.1h transformed with pJBN11B |
| T1013.1h | *A, ridΔ, mus-52Δ, csr-1Δ, his-3+::FOC100X* | T798.1 transformed with pFOC100X |
| T1033.3h | *A, ridΔ, mus-52Δ, csr-1Δ, his-3+::FOC100X, sad-6Δ* | T1013.1h transformed with pJBN1 |
| T1033.8h | *A, ridΔ, mus-52Δ, csr-1Δ, his-3+::FOC100X, sad-6Δ* | T1013.1h transformed with pJBN1 |
| T1035.1h | *a, set-7Δ, ridΔ, mus-52Δ, his-3+::tetO, h4v-flag, chromo1-1(JCT1E)* | T961.5h transformed with pJCT1E |
| T1035.2h | *a, set-7Δ, ridΔ, mus-52Δ, his-3+::tetO, h4v-flag, chromo1-1(JCT1E)* | T961.5h transformed with pJCT1E |
| T1037.3h | *a, rid+ (FOC148C), mus-52Δ, csr-1Δ, his-3+::csr-1+, sad-6Δ* | T996.5h transformed with pFOC148C |

**Table S3.** Antibodies used in this study.

| Epitope | Catalog number | Supplier |
| --- | --- | --- |
| FLAG | F1804 | Sigma |
| GFP | AB290 | Abcam |
| H3 | AB1791 | Abcam |
| H3K9me3 | AB_2532132 | Active Motif |
| HA | 3F10 | Roche |
| Rpb1-S5P | AB5131 | Abcam |

**Table S4.**
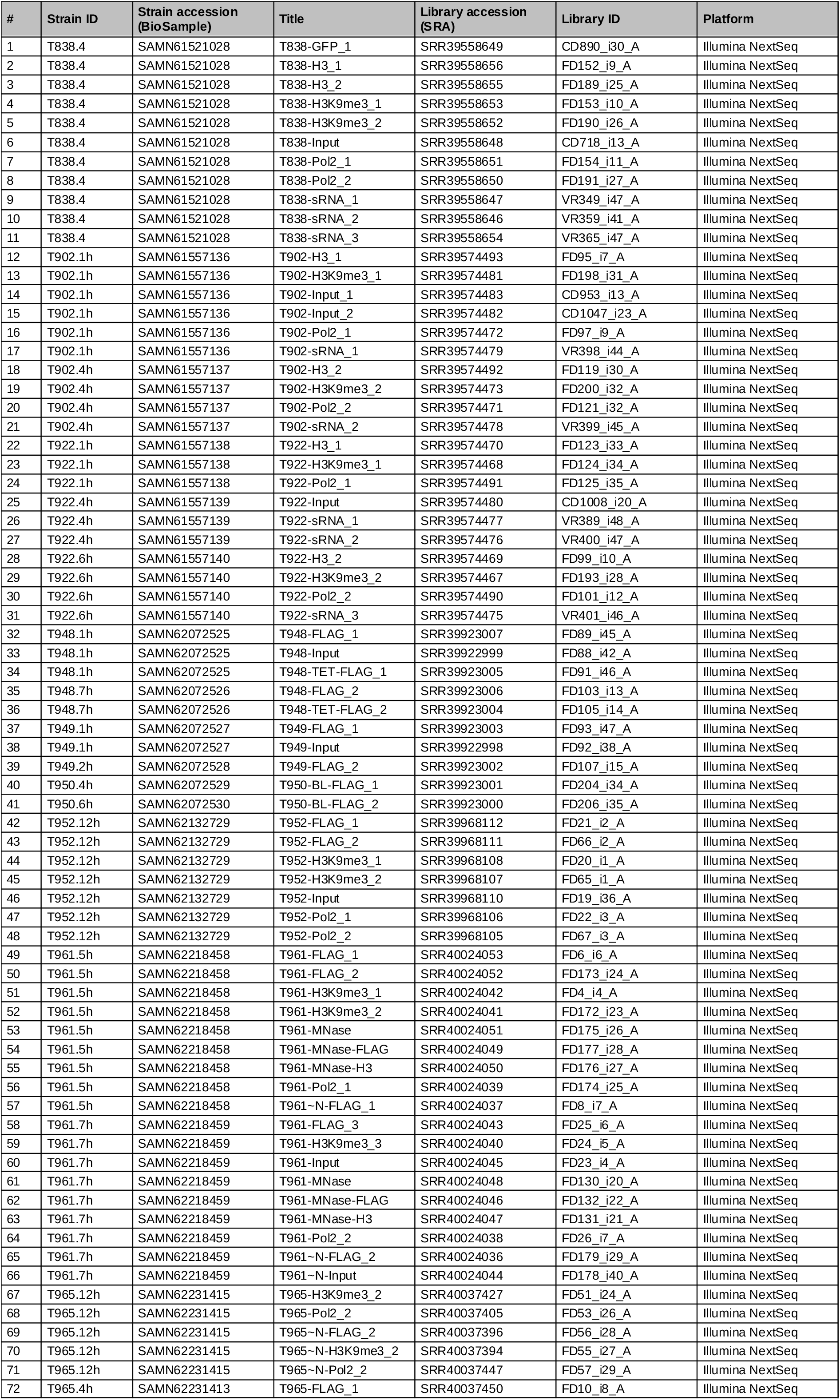
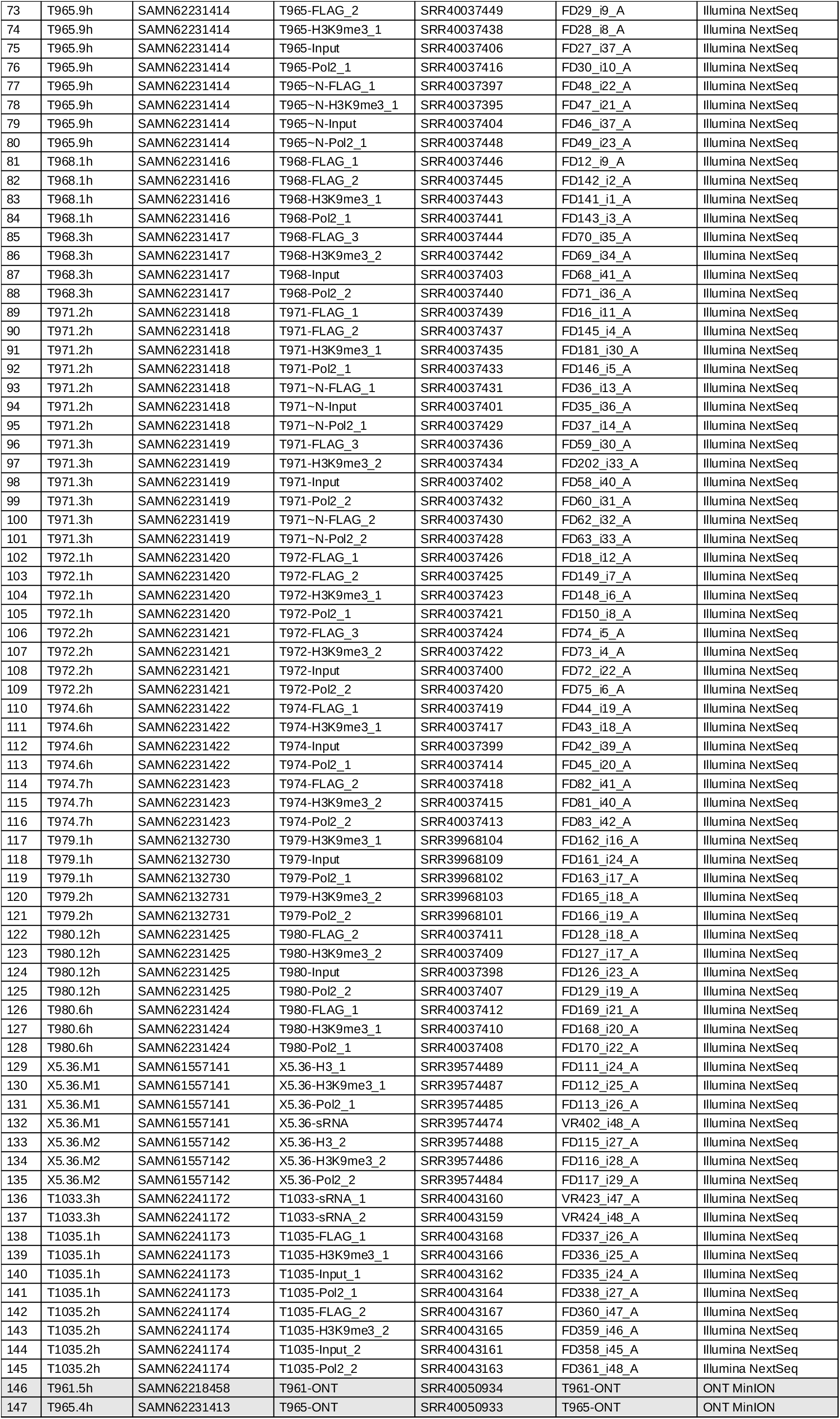
Next-generation sequencing libraries generated and analyzed in this study (ordered by strain).

**Table S5.** Crosses analyzed for RIP in this study.

| Cross ID | Female parent | Male parent | Cross genotype |  |  |  |
| --- | --- | --- | --- | --- | --- | --- |
| FCE1 | T804.8h | T805.1h | *his-3+::csr-1+/his-3+::FOC100T* | *ridΔ/Δ* | *dim-2+/+* |  |
| FCE3 | T804.8h | T805.1h | *his-3+::csr-1+/his-3+::FOC100T* | *ridΔ/Δ* | *dim-2+/+* |  |
| FCE4 | T804.8h | T805.1h | *his-3+::csr-1+/his-3+::FOC100T* | *ridΔ/Δ* | *dim-2+/+* |  |
| FCE5 | T996.5h | T1000.2h | *his-3+::csr-1+/his-3+::FOC100T* | *ridΔ/Δ* | *dim-2+/+* | *sad-6Δ/Δ* |
| FCE6 | T996.8h | T1000.12h | *his-3+::csr-1+/his-3+::FOC100T* | *ridΔ/Δ* | *dim-2+/+* | *sad-6Δ/Δ* |
| FCE7 | T997.7h | T1001.3h | *his-3+::csr-1+/his-3+::FOC100T* | *ridΔ/Δ* | *dim-2+/+* | *h4vΔ/Δ* |
| FCE8 | T997.8h | T1001.4h | *his-3+::csr-1+/his-3+::FOC100T* | *ridΔ/Δ* | *dim-2+/+* | *h4vΔ/Δ* |
| FCE9 | T998.4h | T1002.2h | *his-3+::csr-1+/his-3+::FOC100T* | *ridΔ/Δ* | *dim-2Δ/Δ* |  |
| FCE10 | T998.6h | T1002.5h | *his-3+::csr-1+/his-3+::FOC100T* | *ridΔ/Δ* | *dim-2Δ/Δ* |  |
| FCE24 | T996.5h | T1033.3h | *his-3+::csr-1+/his-3+::FOC100X* | *ridΔ/Δ* | *dim-2+/+* | *sad-6Δ/Δ* |
| FCE25 | T996.5h | T1033.3h | *his-3+::csr-1+/his-3+::FOC100X* | *ridΔ/Δ* | *dim-2+/+* | *sad-6Δ/Δ* |
| FCE26 | T996.5h | T1033.8h | *his-3+::csr-1+/his-3+::FOC100X* | *ridΔ/Δ* | *dim-2+/+* | *sad-6Δ/Δ* |
| FCE27 | T996.5h | T1033.3h | *his-3+::csr-1+/his-3+::FOC100X* | *ridΔ/Δ* | *dim-2+/+* | *sad-6Δ/Δ* |
| FCE28 | T996.5h | T1033.8h | *his-3+::csr-1+/his-3+::FOC100X* | *ridΔ/Δ* | *dim-2+/+* | *sad-6Δ/Δ* |
| FCE29 | T1037.3h | T1033.3h | *his-3+::csr-1+/his-3+::FOC100X* | *rid+/Δ* | *dim-2+/+* | *sad-6Δ/Δ* |
| FCE30 | T1037.4h | T1033.8h | *his-3+::csr-1+/his-3+::FOC100X* | *rid+/Δ* | *dim-2+/+* | *sad-6Δ/Δ* |

## Notes

### Competing Interest Statement

The authors have declared no competing interest.

